# Fragmentation of macrophages during isolation confounds analysis of single cell preparations from mouse hematopoietic tissues

**DOI:** 10.1101/2021.04.28.441876

**Authors:** Susan M Millard, Ostyn Heng, Khatora S Opperman, Anuj Sehgal, Katharine M Irvine, Simranpreet Kaur, Kim M Summers, Cheyenne J Sandrock, Andy C Wu, Graham W Magor, Lena Batoon, Andrew C Perkins, Jacqueline E Noll, Andrew CW Zannettino, David P Sester, Jean-Pierre Levesque, David A Hume, Liza J Raggatt, Allison R Pettit

## Abstract

Mouse hematopoietic tissues contain abundant and heterogeneous populations of tissue-resident macrophages attributed trophic functions in control of immunity, hematopoiesis and bone homeostasis. A systematic strategy to characterise macrophage subsets in mouse bone marrow (BM), spleen and lymph node, unexpectedly revealed macrophage surface marker staining typically emanated from membrane-bound subcellular remnants associated with unrelated cell types. Remnant-restricted macrophage-specific membrane markers, cytoplasmic fluorescent reporters and mRNA were all detected in non-macrophage cell populations including isolated stem and progenitor cells. The profile of macrophage remnant association reflects adhesive interactions between macrophages and other cell types in vivo. Applying this knowledge, reduced macrophage remnant attachment to BM granulocytes in *Siglec1* deficient mice was associated with compromised emergency granulocytosis, revealing a function for *Siglec1*-dependent granulocyte-macrophage interactions. Analysis of published RNA-seq data for purified macrophage and non-macrophage populations indicates that macrophage fragmentation is a general phenomenon that confounds bulk and single cell analysis of disaggregated tissues.

## Introduction

Tissues contain abundant tissue-resident macrophages that contribute to tissue development, homeostasis, regeneration and pathology. Within tissues, sub-populations of resident macrophages are defined based upon location and surface markers (Bleriot et al., 2020; Hume et al., 2019). Organs that have pleotropic function like hematopoietic tissues contain more than one tissue-resident macrophage population with distinct functional contributions to specific biological events. For example, spleen contains CD169^−^F4/80^+^ red pulp macrophages integral in phagocytic clearance of aged/damaged erythrocytes and iron recycling, tingible body macrophages responsible for clearance of apoptotic germinal centre B-cells, CD169^+^F4/80^−^ metallophilic marginal zone macrophages and SIGN-R1^+^F4/80^+^ marginal zone macrophages, both contributing to blood filtration, capturing and presenting/delivering antigens to either white pulp dendritic cells or marginal zone B-cells, respectively (den Haan and Kraal, 2012; Lewis et al., 2019). Similarly, bone and bone marrow (BM) contain abundant F4/80^+^/CD115(CSF1R)^+^ resident macrophages that have been subdivided into bone-associated osteal macrophages (Chang et al., 2008) and those associated with erythroblastic islands (Tay et al., 2020; Yeo et al., 2019) and hematopoietic stem cell (HSC) niches (Chow et al., 2011; Winkler et al., 2010). BM also contains osteoclasts and monocytes and their progenitors, all of which share surface markers with macrophages.

The resident macrophage populations in many tissues have been isolated and profiled by RNA sequencing (RNA-seq) (Summers et al., 2020). However, the cell yields are low compared to their abundance in situ, and it is unclear whether the full diversity of macrophage phenotypes present in vivo have been comprehensively sampled. Mass cytometry analysis across eight murine tissues, including spleen and BM, identified only four tissue-resident macrophage populations. None of these were substantially represented in the BM, where the sum of all macrophage clusters was <1% of the myeloid compartment (Becher et al., 2014). Similarly, high throughput, low coverage single cell RNA-Seq (scRNA-Seq) of BM did not identify a single resident macrophage population (Baccin et al., 2020). High coverage scRNA-Seq of over 40,000 cells across 20 mouse organs to generate a transcriptomic atlas of biology, demonstrated poor alignment between manual macrophage annotation and unbiased whole-transcriptome based clustering (Tabula Muris, 2018).

Conventional macrophage gating strategies in naive mouse tissues indicate macrophages comprise just 0.7% and 1.3% of hematopoietic (CD45^+^) cells in BM and spleen respectively (Cossarizza et al., 2019). These proportions are inconsistent with the abundance of F4/80^+^/CSF1R^+^ resident macrophages in these and other tissues assessed by in situ techniques (Grabert et al., 2020; Sasmono et al., 2003). An estimate of 10-15% of total cells is supported by indirect evidence that macrophages contribute around 10% of total mRNA in most organs in the mouse (Summers and Hume, 2017). Using a positive selection macrophage marker profile (*Csf1r*-EGFP reporter^+^F4/80^+^VCAM-1^+^ coupled with validation using CD169 (Siglec1), ER-HR3, MerTK and/or TIM4 (Kaur et al., 2018)) we improved the efficiency of conventional flow cytometry gating of BM resident macrophages (3.8% of total BM). However, the yield remained low and clear sub-populations reflecting BM macrophage functional diversity were not evident. Herein we took a systematic negative selection approach to identify the distinct functional macrophage subsets in hematopoietic tissue. Our results revealed that macrophages are structurally compromised by processing to obtain single cell suspension, resulting in cell artefacts that have significant implications for interpretation of ex vivo cell analyses.

## Results

### Negative selection to enrich for putative macrophages in hematopoietic tissues

A systematic negative selection strategy was used to identify resident macrophage subsets in hematopoietic/lymphoid organs. Cells isolated from *Csf1r–*EGFP reporter mice (Sasmono et al., 2003) were stained with an antibody panel designed to delineate mature hematopoietic cell lineages and broad progenitor cell subsets (Figure 1A and Figure S1). Defined non-macrophage cell populations (specified in magenta text, Figure 1A) accounted for 91±1% of live cell singlets in BM, 97±2% in spleen and 98±0.4% in lymph node (Table S1). Following exclusion of lineage markers (Ter119, Ly6G, B220, CD4/CD8, NKp46 and CD11c), the remaining cells were sub-gated into three populations: #1, lineage^−^CD11b^+^Ly6G^−^CD115^−/lo^GFP^−/lo^; #2, lineage^−^CD11b^−^Kit^−/lo^GFP^−^; #3, lineage^−^CD11b^−^Kit^−/lo^GFP^+^ (Figure 1A, final 3 panels). Subsets of population #1-3 stained positively for F4/80, VCAM-1 and CD169 (Figure 1B-E). The frequency of F4/80^+^ events within these populations was typically highest in population #1, but also varied between tissues (Table S2). Based on this, the most likely population representing bona fide hematopoietic macrophages is population #1, which represents 5.5±1.4% of total BM cells. This remains much less than the apparent abundance of F4/80^+^ staining observed in BM in situ (Figure 2A-B). Moreover, the population #1 marker profile of CD11b^+^Ly6G^−^CD115^−^ resembles that of neutrophil precursors (Kim et al., 2017; Zhu et al., 2018). Based upon these observations we used imaging flow cytometry to scrutinize the location of the cell surface markers and reporter molecules.

**Figure 1.**
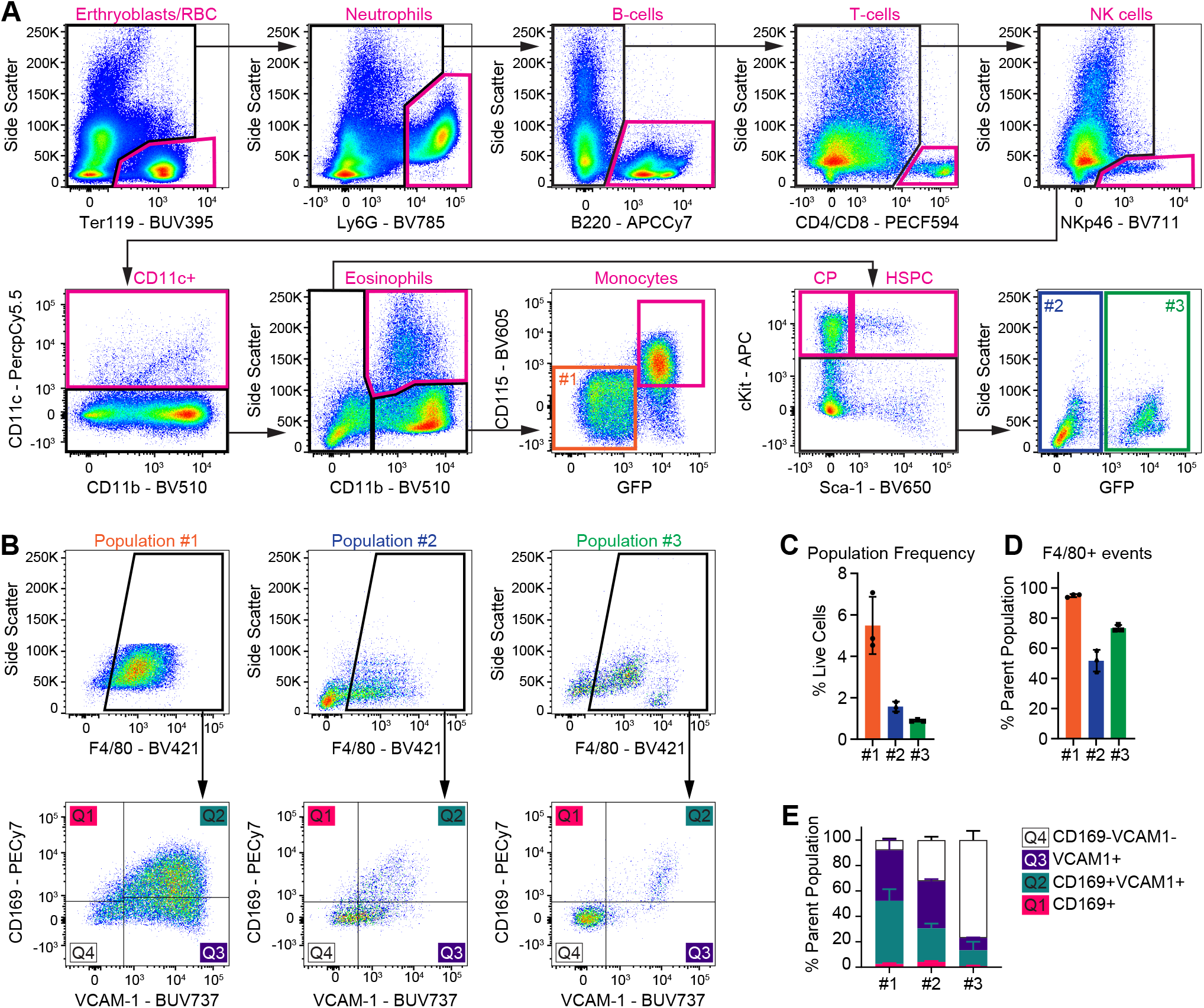
A Negative Selection Approach Identifies Three Putative BM Macrophage Populations. (A) Flow cytometry gating strategy employed to identify and exclude defined non-macrophage hematopoietic populations in BM from *Csf1r*-EGFP reporter mice. Excluded cell populations are labelled in magenta, and remaining putative macrophage populations labelled in orange (#1), blue (#2) and green (3#). Application of this gating strategy to spleen, lymph node and peripheral blood is shown in Figure S1. Cell frequency data for each population in each tissue is summarized in Table S1. (B) Gating strategy employed to examine macrophage marker expression on putative macrophage populations in BM. (C) Frequency of putative macrophage populations in BM. Data are represented as mean ± s.d.; *n* = 3. (D) Frequency of F4/80^+^ staining on putative macrophage populations in BM. These data are summarized in Table S2. (E) Subsetting of F4/80^+^ events based on CD169 and VCAM-1 staining on each of the putative macrophage populations. Data are represented as mean ± s.d.; *n* = 3.

**Figure 2.**
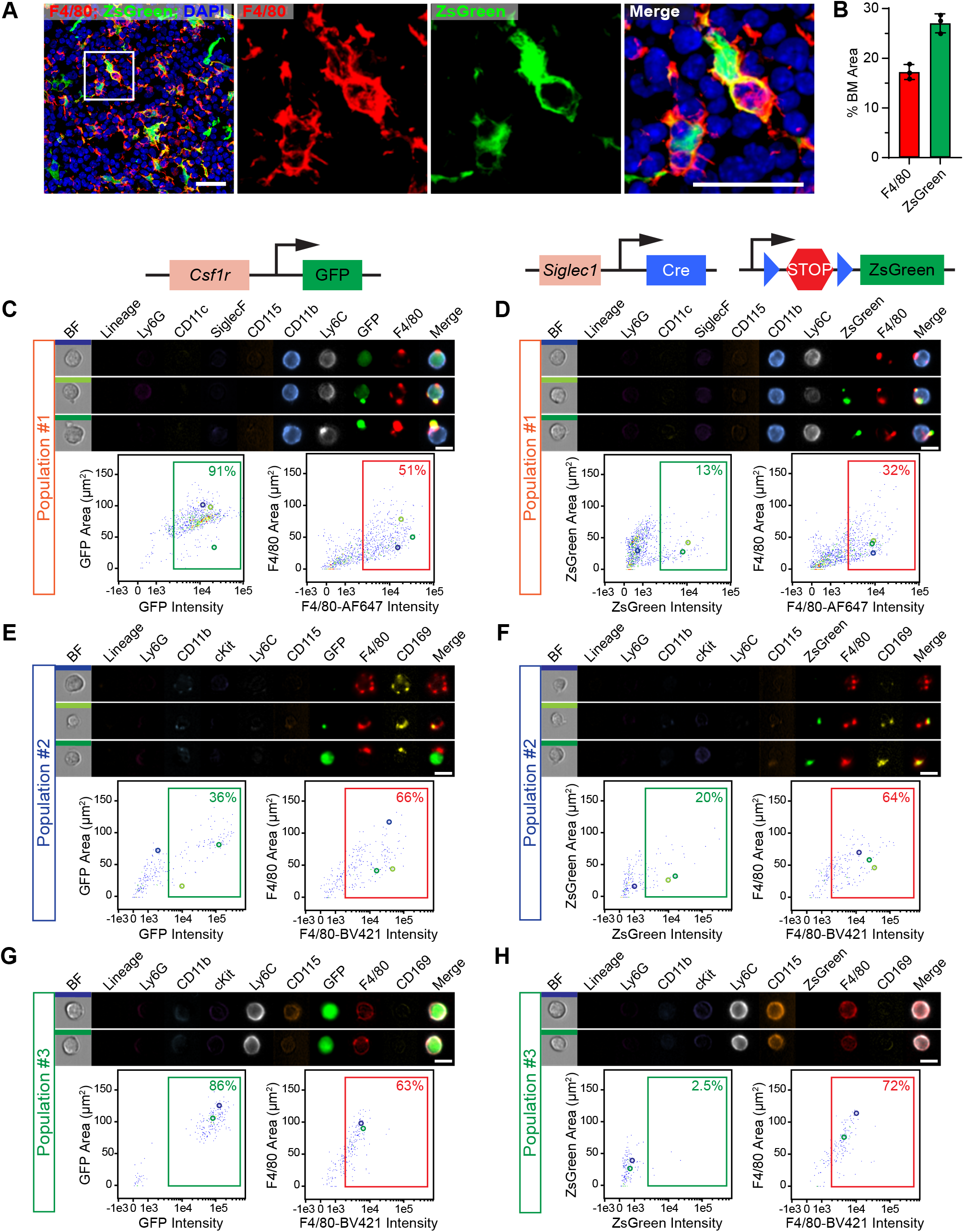
Imaging Flow Cytometry Revealed Macrophage Markers on BM cells in Suspension Are Not Indicative of Intact Macrophages. (A) Co-expression of the cell-surface pan-macrophage marker F4/80 and cytoplasmic ZsGreen reporter in BM from *Siglec1*^Cre^;*R26*^ZsGreen^ reporter mice. Scale bars, 25 μm. (B) Estimation of macrophage frequency in BM by quantification of stain area of the cell-surface marker, F4/80, and the cytoplasmic marker, ZsGreen, in bone cryosections as represented in (A). Data are represented as mean ± s.d.; *n* = 3. (C and D) Representative imaging flow cytometry images from putative macrophage population #1 in BM from *Csf1r*-EGFP (C) and *Siglec1*^Cre^;*R26*^ZsGreen^ (D) mice. The gating strategy is outlined in Figure S2A. The coloured bars on the brightfield (BF) images correspond to the coloured circles on matched plots of stain area against stain intensity, placing the visualized images in the context of the data-set. Scale bar, 10 μm. (E and F) Representative imaging flow cytometry images from putative macrophage population #2 in BM from *Csf1r*-EGFP (E) and *Siglec1*^Cre^;*R26*^ZsGreen^ (F) mice. The gating strategy is outlined in Figure S2D. Scale bar, 10 μm. (G and H) Representative imaging flow cytometry images from putative macrophage population #3 in BM from *Csf1r*-EGFP (G) and *Siglec1*^Cre^;*R26*^ZsGreen^ (H) mice. The gating strategy is outlined in Figure S2D. Scale bar, 10 μm.

### Siglec1^Cre^;R26^ZsGreen^ reporter displays higher fidelity for BM macrophages than Csf1r-EGFP

Reflecting the expression of *Csf1r* mRNA, *Csf1r*-EGFP reporter expression was confirmed in B-cells, neutrophils and monocytes (Figure S2A and S2B; (Sasmono et al., 2003)). SiglecF^+^ eosinophils showed variable expression of the *Csf1r–*EGFP reporter, but no cell surface CD115 (Figure S2A and S2B; (Hawley et al., 2018)). Of these populations, cell surface F4/80 expression was observed on monocytes and eosinophils (Fukushima et al., 2010; Hamann et al., 2007). We also employed a *Siglec1*^Cre^;*R26*^ZsGreen^ conditional reporter for improved specificity. In the BM, in situ imaging showed coincident expression of ZsGreen in F4/80^+^ macrophages (Figure 2A-B). Imaging flow cytometry confirmed ZsGreen reporter was absent from erythroblasts, B-cells and T-cells (included in lineage), neutrophils, eosinophils and monocytes (Figure S2A and S2C), validating its utility as a high fidelity reporter for BM macrophages.

Figure 2C-H shows imaging flow cytometry of the three populations of negatively-selected putative BM macrophages from both *Csf1r*-EGFP and *Siglec1*^Cre^;*R26*^ZsGreen^ mice (gating strategies outlined in Figure S2A and S2D). In *Csf1r*-EGFP mice, population #1 was 91% GFP^+^ with the majority having cytoplasmic distribution consistent with endogenous GFP reporter expression (Figure 2C; median GFP stain area = 76 μm^2^/cell). The same cells expressed surface CD11b, consistent with their being myeloid lineage cells. However, F4/80 staining was polarised, being restricted to 1 or more foci associated with the imaged cell (Figure 2C; median F4/80 stain area = 41 μm^2^/cell). In some instances, GFP expression colocalised with the intense foci of F4/80 expression (Figure 2C). Few cells in this same population expressed cytoplasmic ZsGreen in the *Siglec1*^Cre^;*R26*^ZsGreen^ mice (Figure 2D). When present, it was always coincident with foci of F4/80 (Figure 2D). When antibodies against macrophage markers F4/80, CD169 and VCAM-1 were examined simultaneously they shared the same polarized, focal staining pattern on CD11b^+^ cells (Figure S3A).

Amongst population #2, fewer events expressed *Csf1r*-EGFP reporter than F4/80 and when present GFP distribution was often, but not always, coincident with F4/80 (Figure 2E). In this population *Siglec1*^Cre^;*R26*^ZsGreen^ reporter, when present, was consistently coincident with focal F4/80 staining (Figure 2F). Population #3 cells were brightly GFP^+^ in *Csf1r*-EGFP mice with staining area indicative of large cells (Figure 2G; median GFP stain area = 98 μm^2^). They frequently expressed surface CD115 and also displayed uniform cell surface F4/80 staining (Figure 2G and 2H). However, neither cell surface CD169 nor ZsGreen was detected, indicating this minor ex vivo population does not represent the dominant F4/80^+^ZsGreen^+^ macrophages observed in bone tissue sections (Figure 2A).

In summary, BM macrophages cannot be identified as a distinct population within ex vivo single cell suspensions. Specifically, imaging flow cytometry profiles of BM demonstrate that when unrelated cell types are excluded based upon surface markers, the major residual population was comprised of immature myeloid cells cloaked with fragmented cell remnants likely donated by ramified macrophages. Importantly, these putative macrophage remnants retain cytoplasmic reporter proteins (GFP, ZsGreen).

### Macrophage remnants are found on many distinct cell types and have tissue specific distribution patterns

The presence of macrophage remnants was not restricted to immature myeloid populations enriched by negative selection. Figure 3 shows that macrophage remnants were associated with mature hematopoietic lineage cells, including erythroblasts, B-cells, T-cells and neutrophils, in both BM (Figure 3A, gated as outline in Figure S3B and S3C) and spleen (Figure 3B, gated as outlined in Figure S3D and S3E). With the exception of monocytes, which displayed endogenous cell-surface F4/80 expression, the F4/80 stain area for each cell type was significantly lower than the stain area of a defining cell surface marker (Figure 3C and 3D).

**Figure 3.**
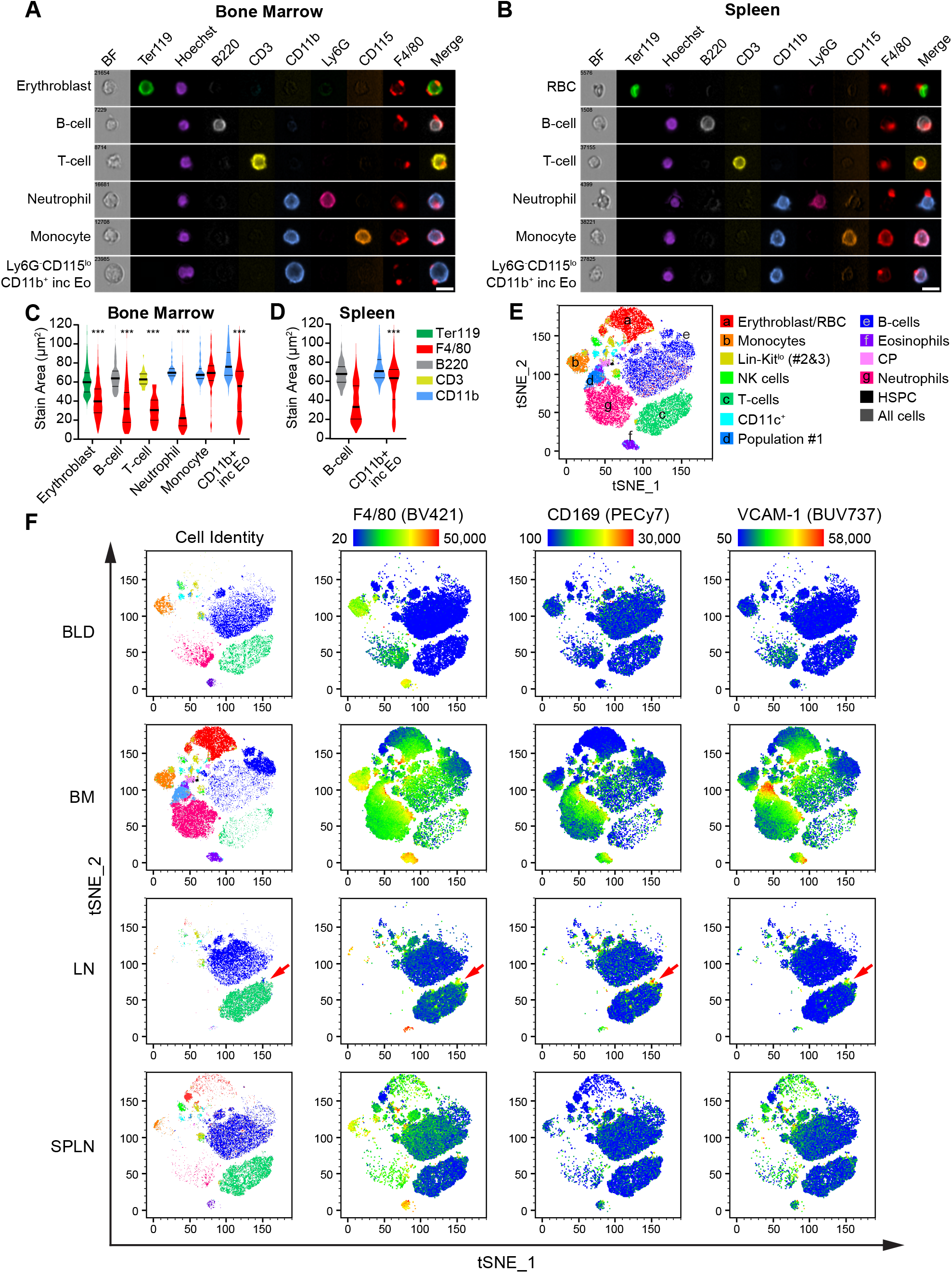
Macrophage Markers Are Detected Across Many Cell Lineages in Hematopoietic Tissue-Derived Cell Suspensions. (A and B) Representative images of F4/80 staining associated with hematopoietic cell lineages that are not macrophages, as visualized in bone marrow (A) and spleen (B) using imaging flow cytometry according to the gating strategy outlined in Figure S3. The merge image consists of F4/80 with a relevant cell-surface marker (Ter119, erythroblasts; B220, B-cells; CD3ε, T-cells; CD11b, neutrophils, monocytes, Ly6G^−^CD115^lo/-^ CD11b^+^ cells; this final population includes eosinophils). (C and D) Paired comparison of stain area for a defining cell-surface marker compared to F4/80 for the cell populations displayed in A and B. Data displayed as violin plots with median and 25^th^/75^th^ percentiles. Kruskal-Wallis test with Dunn’s correction for multiple comparisons; *** p<0.0001. n = 26-1086 for each cell type. Too few F4/80^+^ splenic T-cells, neutrophils and monocytes were captured in the imaging flow cytometry assay for statistical analysis. (E) Mapping of hematopoietic cell lineages in conventional flow cytometry data pooled from peripheral blood, BM, lymph node (LN) and spleen (SPLN) to a t-SNE plot. Indicated cell types are gated as outlined in Figure 1A and S1. (F) For each tissue, the tSNE plot is used to display cell composition (first column, color-coded according to (E)) and a heatmap of staining intensity for the macrophage-expressed surface markers F4/80, CD169 and VCAM-1 (subsequent columns). A hot-spot of macrophage marker staining on LN T-cells is indicated (red arrows).

To visualize the extent to which macrophage remnants were present across hematopoietic lineages in single cell suspensions, conventional flow cytometry data from BM, spleen, lymph node and blood (Figure 1A and Figure S1) were pooled and dimensionality reduction (t-SNE) applied based on lineage markers (excluding GFP, F4/80, CD169 & VCAM-1) (Figure 3E). Each tissue was then assessed independently (Figure 3F) with the first column showing the cellular composition of the tissue, and subsequent columns a heat-map representation of the staining intensity for the macrophage markers F4/80, CD169 and VCAM-1. In addition to the global overview of macrophage marker distribution provided by the tSNE heatmaps, the frequency of macrophage marker staining on each defined cell type (gated as outlined in Figure 1) is shown in Figure S4. Peripheral blood was analysed as a comparator obtained without mechanical disruption. In blood, F4/80 staining was restricted to cells with confirmed endogenous cell-surface F4/80 expression and, as expected, CD169 and VCAM-1 were absent (Figure 3F and Figure S4D - S4G). In BM, subsets of all defined cell types had detectable F4/80, CD169 and VCAM-1 (Figure 3F), consistent with imaging flow cytometry showing macrophage remnants (Figure 3A). Macrophage marker promiscuity varied between tissues and between cell populations within tissues. For example, within lymph node, a portion of T-cells had high intensity staining for F4/80, CD169 and VCAM-1 (Figure 3F).

In spleen, in addition to high intensity F4/80 staining on cells with endogenous F4/80 expression (monocytes and eosinophils), many B-cells had moderate intensity F4/80 staining (Figure 3F) whereas CD169 and VCAM-1 staining were more restricted (Figure 3F). In contrast, minimal macrophage marker stain was observed on T-cells. This more selective pattern of remnant attachment raised the possibility that macrophage remnant binding is not random. In spleen F4/80^+^CD169^−^ macrophages are restricted to the red pulp, whereas CD169^+^F4/80^−^ metallophilic marginal zone macrophages are situated at the border of B-cell follicles and the marginal zone (Idoyaga et al., 2009) (Figure 4A). Within the red pulp there is a higher density of B-cells than T-cells (Figure 4A) and accordingly, in situ quantification showed a higher proportion of B220^+^ stain within close proximity of F4/80^+^ stain compared to CD3 stain (Figure 4A and 4B). Similarly, very few T-cells were observed in the marginal zone with a higher proportion of B220^+^ stain within close proximity of CD169^+^ stain (Figure 4A and 4C). The distribution frequency of F4/80 and CD169 staining on B-cells and T-cells by flow cytometry (Figure 4D and 4E) followed a similar pattern, with more F4/80^+^ events (Figure 4F) and more CD169^+^F4/80^−^ events (Figure 4G) observed within the B-cell gate than the T-cell gate. Hence the macrophage remnant profile on splenic lymphocytes correlated with their proximity to specific macrophage populations in vivo. This suggests that macrophage remnants preferentially adhere to cells with which they were interacting prior to mechanical disruption.

**Figure 4.**
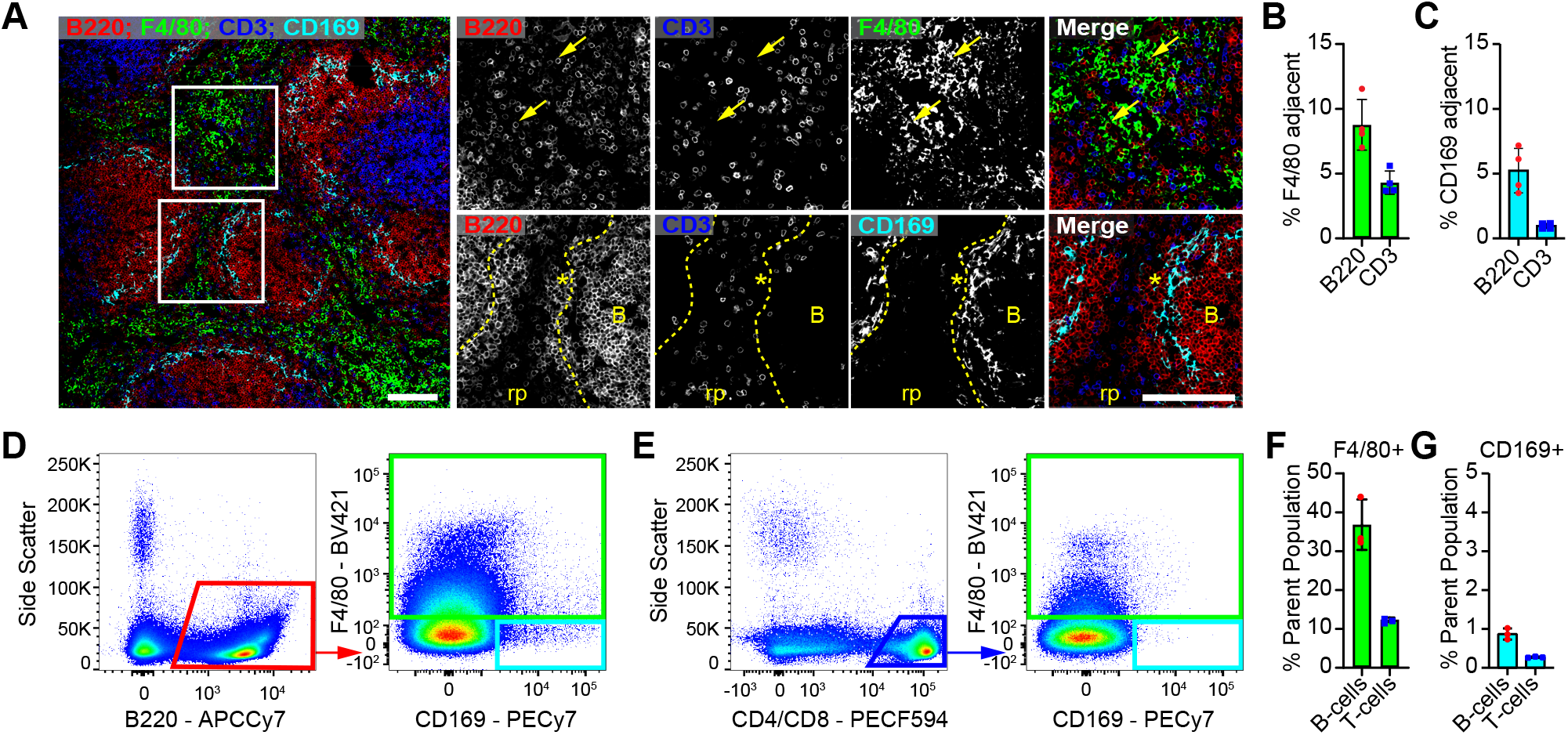
Extent of Lymphocyte Colocalization with Splenic Macrophage Subsets in situ is Consistent with Promiscuous Macrophage Marker Staining on Dissociated Splenic Lymphocytes. (A) Cryosections of C57BL/6J mouse spleen stained for B-cells (B220), T-cells (CD3), red pulp macrophages (F4/80) and metallophilic marginal zone macrophages (CD169). The top panel of images highlight a region of red pulp where B-cells are more populous than T-cells and more likely to be present in areas with intense F4/80 staining (yellow arrows). The bottom panel images highlight a region where CD169^+^ macrophages demarcate the boundary between the B-cell follicle (B) and marginal zone (*). In this image T-cells are mostly restricted to the small area of red pulp (rp) visible between the follicles. Scale bars, 100 μm. (B) Assessment of the colocalization of B-cells and T-cells with red pulp macrophages by quantitation of the frequency of B220 or CD3 staining adjacent to F4/80 staining (within 3 pixels). Data are represented as mean ± s.d*.; n =* 4. (C) Assessment of the colocalization of B-cells and T-cells with metallophilic marginal zone macrophages by quantitation of the frequency of B220 or CD3 staining adjacent to CD169 staining (within 3 pixels). Data are represented as mean ± s.d*.; n =* 4. (D and E) Gating strategy used to assess the frequency of B-cells (D) and T-cells (E) that were either F4/80^+^ or CD169^+^F4/80^−^. Data was pre-gated as shown in Figure S1A. (F and G) Frequency of B-cells and T-cells that are either F4/80^+^ (F) or CD169^+^F4/80^−^ (G) by flow cytometric analysis. Data are represented as mean ± s.d.; *n* = 3.

### Macrophage remnants include cytoplasmic components

In BM isolated from *Siglec1*^Cre^;*R26*^ZsGreen^ animals, 87±2% of CD45^+^F4/80^+^ events were classified by visual inspection as CD45^+^ cells with associated F4/80^+^ subcellular remnants. Significantly, 52±7% of these cells had detectable cytoplasmic ZsGreen reporter colocalized with at least one attached remnant (Figure 5A). Figure 5B shows confocal imaging of an F4/80^+^ remnant attached to a B220^+^ B-cell, within freshly isolated BM. F4/80 and B220 staining are clearly present on the membrane of the subcellular remnant and intact cell, respectively, and ZsGreen fluorescence was encapsulated within F4/80^+^ membrane. The imaged subcellular remnant is 1-2 μm in diameter, making it substantially larger than the typical 50-500 nm extracellular microvesicle (van Niel et al., 2018). To determine whether macrophage mRNA is retained within remnants and contributes to bulk cell sorted expression profiles, we isolated hematopoietic stem and progenitor cell (HSPC) and committed progenitor (CP) cell populations. These cell populations were chosen as they do not natively express mature macrophage genes and macrophages influence their niche in vivo via direct cell interactions (Hur et al., 2016). Both imaging (Figure 5C) and conventional (Figure 5D and 5E) flow cytometry analysis of CD117/Kit^+^ enriched BM confirmed attachment of F4/80^+^ macrophage remnants containing ZsGreen cytoplasmic reporter on all HSPC subsets (gated as outlined in Figure S5A and S5B). Sorted CP (sub-fractioned based on F4/80 and ZsGreen staining) and total HSPC all demonstrated equivalent expression levels of the stem cell marker, *Kit* (Figure 5F and 5G). Macrophage-specific marker transcripts were readily detectable in total HSPC and were present in F4/80+ CP populations at comparable levels to in vitro differentiated BM-derived macrophages (Figure 5F and 5H). Exclusion of F4/80^+^ events from the CP population eliminated detection of not only *Adgre1* (which encodes F4/80) but also mRNA for macrophage markers *Aif1, Cd68* and *Mertk* (Figure 5F and 5H). Conversely, exclusion of ZsGreen^+^ events from sorted F4/80^+^ CP did not impact detection of macrophage-specific genes, indicating mRNA detection sensitivity exceeded fluorescent reporter detection in this assay.

**Figure 5.**
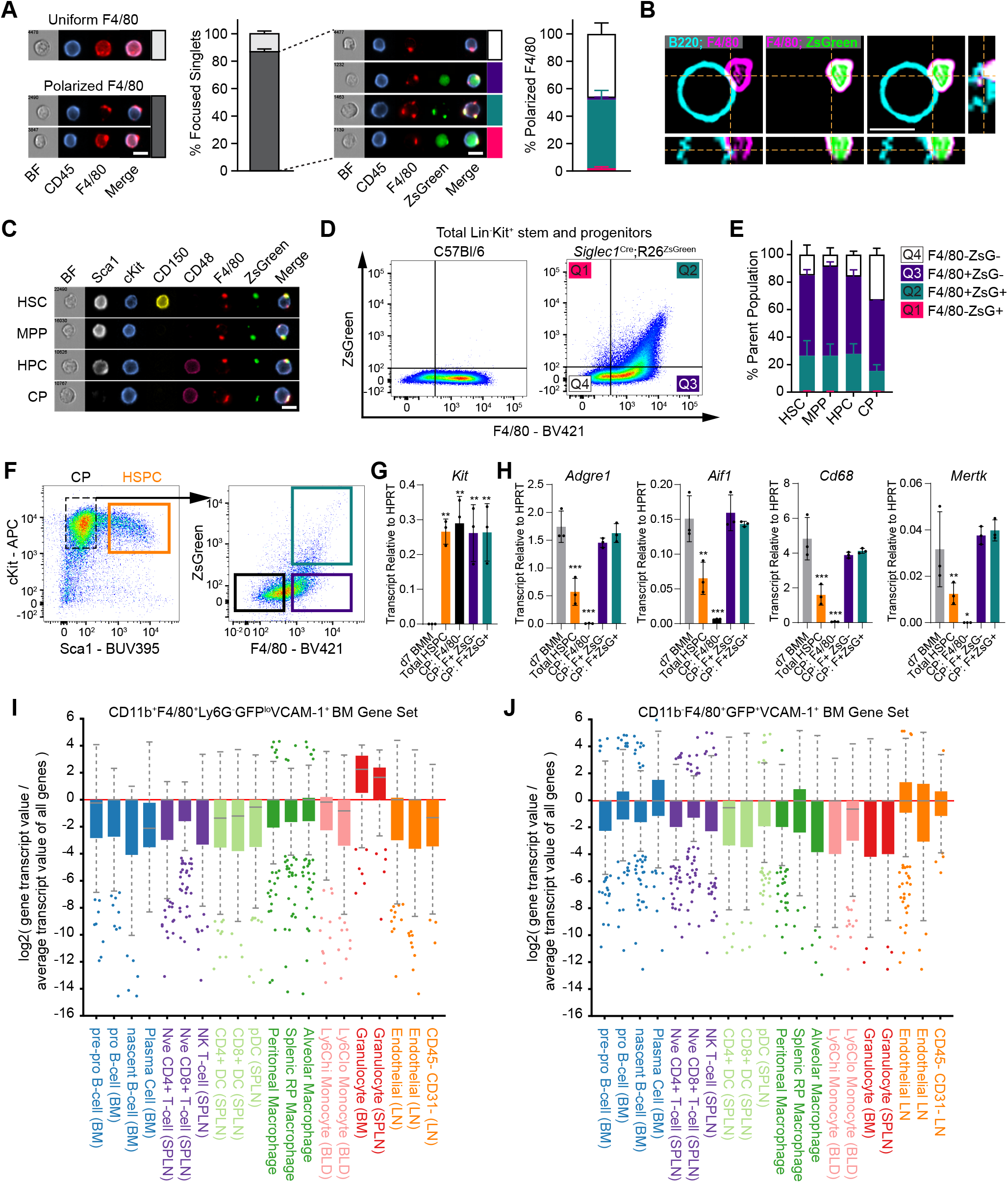
Macrophage Remnants Contain Cytoplasm, Including Robustly Detectable mRNA. (A) BM isolated from *Siglec1*^Cre^;*R26*^ZsGreen^ mice was visualized by imaging flow cytometry. CD45^+^F4/80^+^ events were manually categorized by whether F4/80 staining was polarized or uniformly, and exclusively, cell surface. Polarized events were further categorized based on whether ZsGreen was observed in the primary cell event and/or associated with F4/80 remnant staining. Scale bar, 10 μm. Graphical data are represented as mean ± s.d.; *n* = 4. (B) Confocal imaging of a *Siglec1*^Cre^;*R26*^ZsGreen^ B-cell in BM cell suspension stained for F4/80 (magenta) and B220 (cyan), shown in orthangonal view. Scale bar, 5 μm. (C) F4/80^+^ZsGreen^+^ macrophage remnants visualized on all HSPC subsets in Kit-enriched BM from *Siglec1*^Cre^;*R26*^ZsGreen^ mice. HSPC subsets (CD48^+^ hematopoietic progenitor cell (HPC), CD48^−^CD150^−^ multipotent progenitor (MPP) and CD48^−^CD150^+^ hematopoietic stem cell (HSC)) were gated as outlined in Figure S5A. (D and E) Frequency of F4/80^+^ZsGreen^+^ staining on HSPC subsets enumerated by conventional flow cytometry. The quadrant gates shown for total Lin^−^Kit^+^ cells (D) were applied to each HSPC subset, gated as outlined in Figure S5B, and graphed in (E). Data are represented as mean ± s.d*.; n =* 3. (F – H) Kit-enriched BM pooled from *Siglec1*^Cre^;*R26*^ZsGreen^ mice was sorted into four populations for subsequent RNA expression analysis. Expression of the HSPC marker, *Kit* (G), and macrophage expressed markers (H; *Adgre1*, *Aif1*, *Cd68*, *Mertk*) in the HSPC and CP subsets shown in (F) as compared to in vitro differentiated BM macrophages (d7 BMM). Data are represented as mean ± s.d*.; n =* 3 independent sorts. One-way ANOVA with Dunnett correction for multiple comparisons; * p=0.046, ** p<0.005, *** p<0.0001. (I and J) CD11b^+^ (I) and CD11b^−^ (J) BM populations were sorted based on positive macrophage marker selection, as outlined in Figure S5C, and subjected to RNA-Seq analysis. The prevalence of the top 200 most abundant transcripts from each population is displayed across a subset of the ImmGen ULI RNA-seq data-set (GSE127267). Similar analysis for a CD11b^+^ BM population sorted by an alternate strategy is shown in Figure S5D.

We examined RNA-Seq data from BM cells sorted from *Csf1r*-EGFP mice by positive selection for macrophage markers (gated as outlined in Figure S5C). The most abundant transcripts in these positively selected populations were not enriched within the ImmGen ULI-RNA-Seq monocyte/macrophage datasets (GSE127267). Rather, the CD11b^+^ ‘macrophage’ population (CD11b^+^F4/80^+^Ly6G^−^GFP^lo^VCAM-1^+^) displayed a granulocyte signature and the CD11b^−^ ‘macrophage’ population (CD11b^−^F4/80^+^GFP^+^VCAM1^+^) showed weak similarity with plasma cells and lymph node endothelial cells (Figure 5I and 5J). An alternate sorting strategy for CD11b^+^ ‘macrophages’ (CD11b^+^F480^+^CD169^+^CX3CR1^−^) without Ly6G negative selection also showed a clear granulocyte signature (Figure S5D). In publicly available datasets (Gonzalez et al., 2017; Li et al., 2019; Mildner et al., 2017) curated previously (Summers et al., 2020) and analysed independently, positively selected macrophage populations frequently had granulocyte markers at higher abundance than macrophage markers (Figure S6A and S6B). Erythroblast transcripts were also detected in these populations (Figure S6C).

### BM F4/80^+^Ly6G^+^ events exemplify the macrophage origin of F4/80^+^ remnants

F4/80^+^Ly6G^+^ events detected by traditional flow cytometry have previously been identified and purified as erythroblastic island macrophages (EIM), also expressing VCAM-1, CD169, and ER-HR3 (Jacobsen et al., 2014; Li et al., 2019). Imaging flow cytometry indicated that this dual expression is a consequence of macrophage remnant binding to Ly6G^+^ neutrophils (Figure 3A). To confirm that the F4/80 remnant is donated from macrophages, we examined the impact of inducible depletion of all BM macrophage subsets using diphtheria toxin (DT) treatment of *Siglec1*^DTR/+^ mice (Kaur et al., 2018; Miyake et al., 2007). DT treatment eliminated ramified F4/80 high expressing macrophages (Figure 6A). In contrast F4/80^+^CD115^hi^CD169^−^ cells (myeloid progenitors and monocytes) were expanded, not depleted, in this model (Figure 6A, 6B, 6D and 6E). BM neutrophil frequency was also maintained during DT treatment of *Siglec1*^DTR/+^ mice (Figure 6B and 6E), whereas F4/80 co-staining on Ly6G^+^ neutrophils was eliminated (Figure 6C and 6F). Hence, F4/80^+^Ly6G^+^ events clearly represent fragmented F4/80^+^Ly6G^−^ macrophage remnants adhered to F4/80^−^Ly6G^+^ neutrophils.

**Figure 6.**
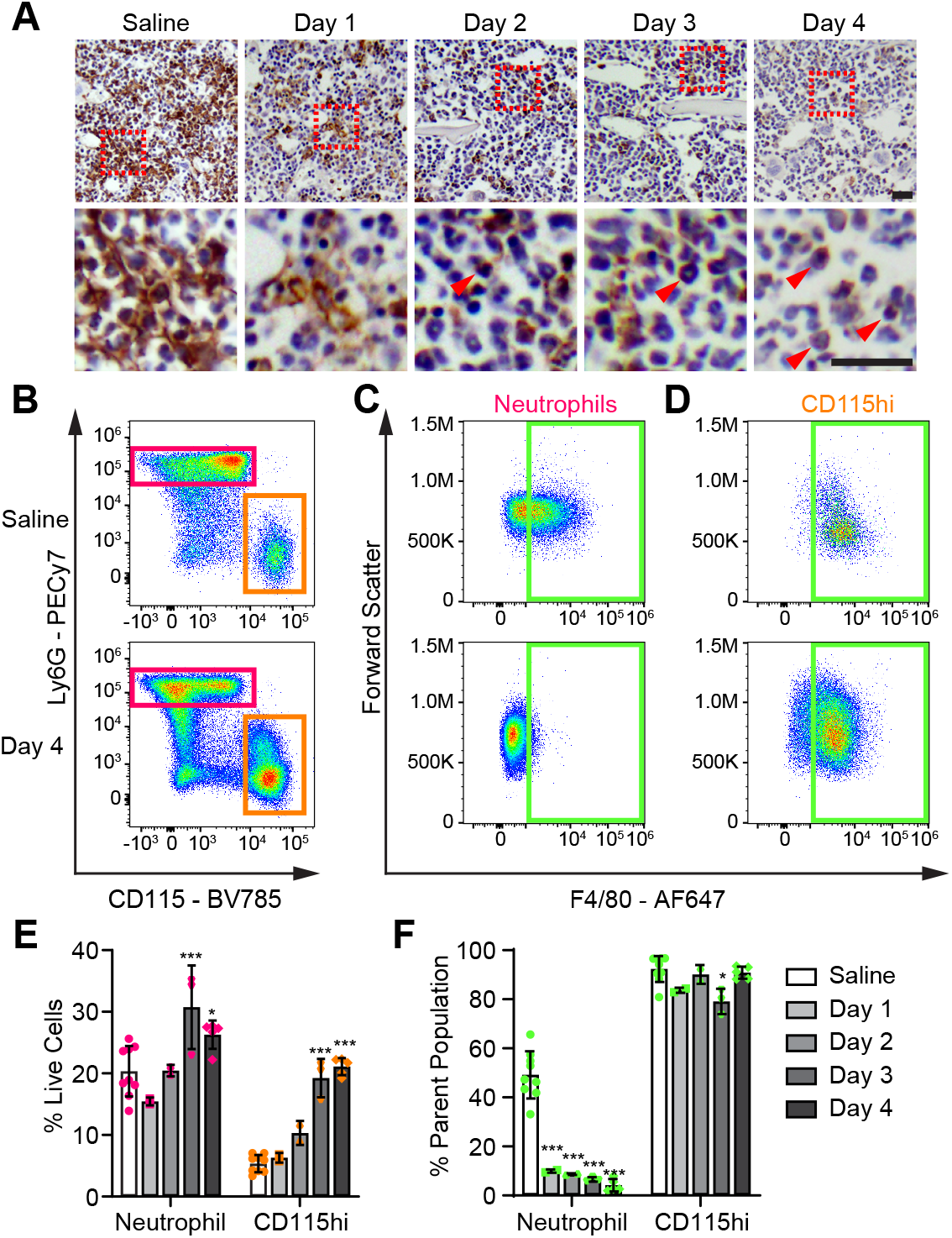
CD169^+^ Macrophage Depletion Eliminates F4/80^+^ Macrophage Remnant Staining. Macrophages were depleted in *Siglec1*^DTR/+^ mice by daily 10 μg/kg diphtheria toxin (DT) injections, starting on Day 0. (A) Immunohistochemical staining of bone marrow (BM) shows a reduction in total F4/80 staining at Day 1 and a complete loss of ramified F4/80^+^ macrophages by Day 4. Small, round F4/80^+^ cells remain evident in the BM following DT treatment (red arrowheads). Scale bars, 25 μm. (B, C and D) Representative flow cytometry plots of BM from saline treated control mice (top panel) and at Day 4 of DT mediated macrophage depletion (bottom panel). Neutrophils (magenta gate) and CD115^hi^ (orange gate) populations were gated from CD11b^+^ BM (B), and F4/80 (green) assessed on these populations (C and D). (E) Minimal impact on neutrophil frequency was observed across the macrophage depletion time course, while CD115^hi^ cell frequency increased. Two-way ANOVA with subsequent comparison to saline control, Tukey correction for multiple comparisons; * p=0.011, *** p<0.0001. (F) Macrophage depletion eliminated F4/80 staining on neutrophils, whereas little impact was observed on F4/80 staining on CD115^hi^ cells. Two-way ANOVA with subsequent comparison to saline control, Tukey correction for multiple comparisons; * p=0.015, *** p<0.0001.

### Knockout of the adhesion molecule CD169 is characterised by selective reduction of F4/80^+^ remnants on granulocytes and HSPC

*Siglec1*^DTR/DTR^ (CD169-KO) mice lack detectable CD169 protein in the BM (Figure 7A) but there is no impact on total F4/80^+^ cells or their distribution (Figure 7A, 7B and 7C). Consistent with the first described CD169-KO model (Oetke et al., 2006), the immune cell composition of the BM was unchanged compared to wild-type mice (Table S3). In wild-type mice, the proportion of CD169^+^ and F4/80^+^ events was similar within HSPC or granulocyte populations (both neutrophils and eosinophils) suggesting remnants attached to these cells had been donated by F4/80^+^CD169^+^ macrophages (Figure S4A). In CD169-KO mice, targeted inspection of HPSC, neutrophil and eosinophil populations showed a selective robust reduction of F4/80^+^ or VCAM-1^+^ remnants, respectively (Figure 7D and 7E). These data indicate that CD169 contributes to the capacity of macrophages to interact with progenitors and other myeloid cells.

**Figure 7.**
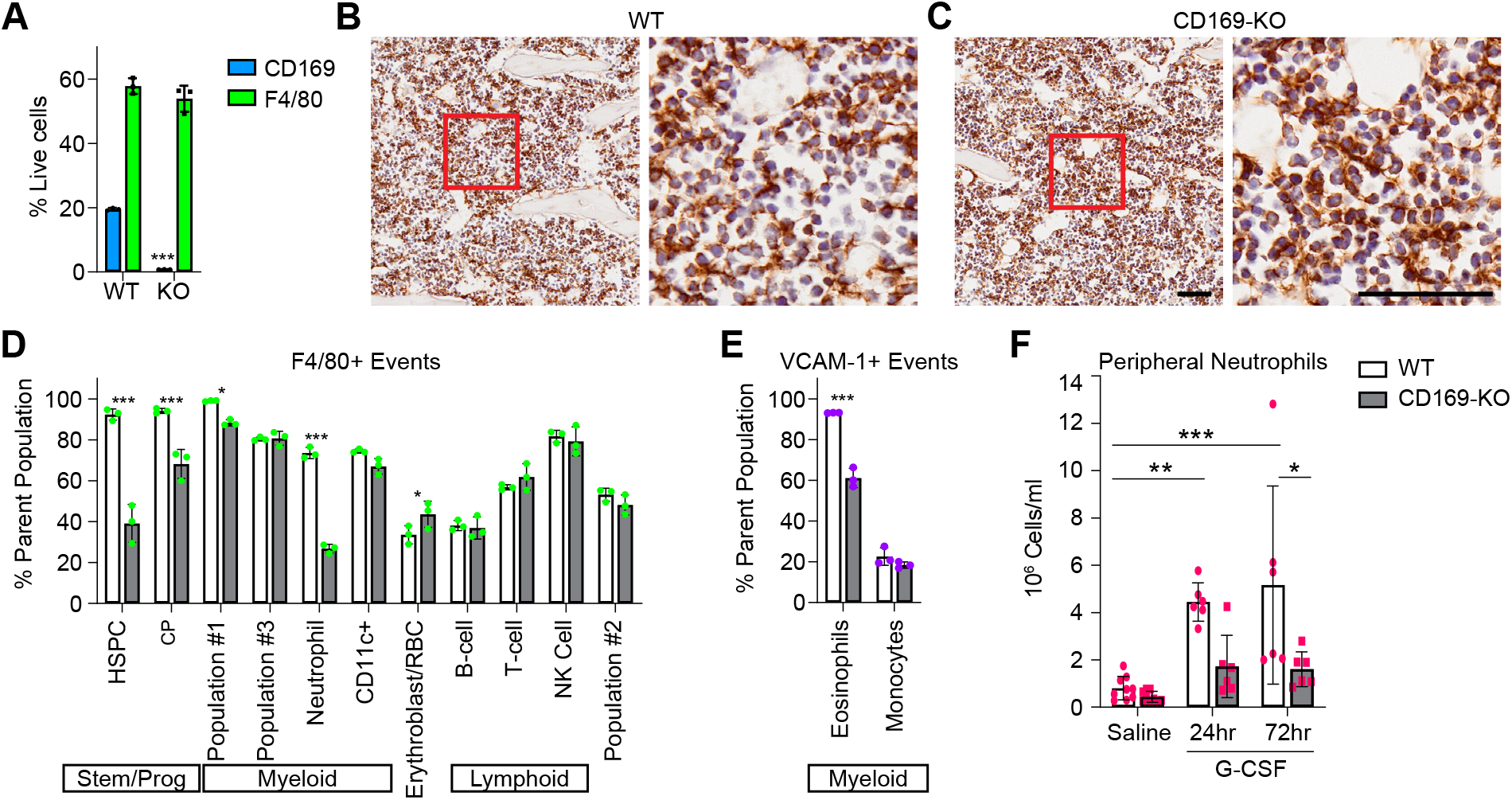
Knockout of *Siglec1* (CD169-KO) Selectively Impacts Macrophage Remnant Adhesion to Granulocytes. (A) CD169-KO eliminates CD169^+^ events, but not F4/80^+^ events, in total BM assessed by conventional flow cytometry. (B and C) Immunohistochemistry for F4/80 in bone isolated from WT (B) and CD169-KO (C) mice. Scale bars, 25 μm. (D and E) CD169-KO selectively impacts the frequency of F4/80^+^ (D) or VCAM-1^+^ (E) macrophage remnants on hematopoietic cell populations in BM. Populations were gated as outlined in Figure 1A. (F) CD169-KO blunted the rapid and sustained increase in peripheral neutrophil numbers in response to bi-daily 125 μg/kg G-CSF treatment, as compared to WT mice. Data are pooled from 4 independent experiments and represented as mean ± s.d*.; n =* 6-9. Two-way ANOVA with Tukey correction for multiple comparisons. * p = 0.011, ** p = 0.003, *** p = 0.0003.

To explore the functional importance of this interaction, we examined granulocyte-colony stimulating factor (G-CSF) induced granulopoiesis in CD169-KO mice. BM and peripheral granulocyte progenitors, precursors and mature granulocytes were gated as per Figure S7A and S7C. The CD169-KO had no effect on BM granulopoiesis induced by G-CSF (Figure S7B). However, in CD169-KO there was a substantial reduction of the peripheral granulocytosis elicited by G-CSF administration (Figure 7F and Figure S7D). This is indicative that suboptimal granulocyte interactions with macrophages compromised their egress from BM to blood.

## Discussion

Our data provide comprehensive evidence that the majority of tissue-resident macrophages are mechanically disrupted during preparation of single cell suspensions from hematopoietic tissues. The negative selection strategy employed to analyse hematopoietic/lymphoid organs accounted for 98.6-99.9% of flow cytometric events in BM, spleen and lymph node without revealing a single resident macrophage subset. Positive staining for macrophage markers was a consequence of other cells being cloaked with remnants. These observations are consistent with the report that subcapsular sinus macrophages are fragmented during lymph node single cell preparation and attached to IL-7 receptor expressing T-cells (Gray et al., 2012). Key observations made herein were that the cell-attached macrophage remnants contained not only membrane and associated proteins at high molecular density, but also intracellular contents including significant amounts of mRNA. This technical phenomenon has contributed to the misassignment of macrophage identity to flow cytometry gated populations in hematopoietic tissues, including our own studies reporting F4/80^+^Ly6G^+^ cell events as EIM (Jacobsen et al., 2014; Kaur et al., 2018). Indeed, Li et al. used a distinct approach to isolate and profile F4/80^+^ EIM based upon an *Epor*-EGFP transgene (Li et al., 2019) but analysis herein of their data revealed strong enrichment for immature myeloid transcripts in the putative EIM. Nonetheless, we showed these ex vivo analyses can still provide valuable and robust biological information. Coupling conventional flow cytometry with in situ approaches (Kaur et al., 2018) or bespoke isolation strategies that maintain multicellular aggregates (Bisht et al., 2020; Seu et al., 2017; Tay et al., 2020; Yeo et al., 2019; Yeo et al., 2016) strengthens data interpretation, and confirms some instances of unconventional co-expression of immune markers (Abtin et al., 2014; Yates et al., 2013).

Macrophage remnants are similar in size to oncosomes, the largest described plasma-membrane derived microvesicles (van Niel et al., 2018). As intact macrophages could not be identified within hematopoietic tissue preparations, it is unlikely that remnants represent extracellular vesicles shed by macrophages in vivo. Rather it appears that the integrity of highly ramified tissue-resident macrophages has been disrupted en masse by the fluid and/or mechanical shear stress inherent in preparing single cell suspensions. Given the absence of an intended vesicular budding/fission event, it is remarkable that macrophage disruption results in structurally stable remnants that retain intracellular contents. Our presumption is that macrophages fail at their structurally narrowest points facilitating membrane fusion due to self-healing capacity of lipid membranes (Sych et al., 2018).

Our results cannot distinguish whether the presence of remnants reveals in vivo cell-cell interactions or reflects attachment of remnants to cell populations with appropriate molecular adhesion profiles during isolation. We favour the former interpretation. Based upon the preferential binding of CD169^+^ macrophage remnants to granulocyte populations in BM we tested and confirmed a role for CD169 in control of granulocyte egress into blood during G-CSF-induced granulopoiesis. The preferential binding of F4/80^+^CD169^−^ macrophage remnants to erythroblasts is more difficult to reconcile with previous evidence (Bisht et al., 2020; Chow et al., 2013; Seu et al., 2017; Tay et al., 2020) but is consistent with in vitro experiments demonstrating that murine CD169 does not bind murine erythroblasts (Kelm et al., 1994).

The disruption of macrophages during tissue disaggregation is not surprising given their remarkable ramification and adhesive properties (Grabert et al., 2020). Kupffer cell disruption and remnant binding to liver endothelial cells has been previously visualized (Lynch et al., 2018). Meta-analysis of bulk sequenced sorted cell populations across 14 different tissues demonstrated that contamination of macrophage data-sets with genes expressed by unrelated cell types is universal (Summers et al., 2020). The reciprocal contamination is also a significant issue, likely leading to false assignment of expression of macrophage-specific genes such as *Csf1r* to non-macrophages. Tissue digestion of spleen did not markedly change the splenic myeloid flow cytometry profile, compared to mechanical disruption (Fujiyama et al., 2019). Enzymatic tissue digestion does not prevent macrophage fragmentation, although it may facilitate isolation of subsets of intact macrophages from some tissues, including liver (Lynch et al., 2018) and lung (Becher et al., 2014; Misharin et al., 2013).

Macrophage fragmentation and remnant binding likely explains why macrophage signatures are poorly represented or challenging to interpret in large scale molecular studies of organ biology (Baccin et al., 2020; Becher et al., 2014; Gautier et al., 2012; Tabula Muris, 2018). It may also reflect why mapping of BM macrophage origin has been conspicuously absent from comprehensive ontogeny studies (Ginhoux and Guilliams, 2016; Liu et al., 2019), and suggests that systematic assessments (Abram et al., 2014; Baccin et al., 2020) of single cell preparations are insufficient for evaluating tissue-resident macrophages. Overall, our observations expose that the current knowledge base is incomplete and probably incorrect with respect to tissue-resident macrophage frequency, diversity and molecular profile.

## Acknowledgements

This work was supported by Mater Foundation, The University of Queensland Postgraduate Scholarship and Mater Research Frank Clair Scholarship (LB), a Veronika Sacco Clinical Cancer Research Fellowship from the Florey Medical Research Foundation, University of Adelaide (JEN), a National Health and Medical Research Council Research Fellowship (JPL, 1136130) and an Australian Research Council Future Fellowship (ARP, FT150100335). This work was carried out at the Translational Research Institute (TRI) which is supported by a grant from the Australian Government. The Translational Research Institute (TRI) Microscopy, Histology and Flow Cytometry Core Facilities contributed technical expertise and the TRI Biological Research Facility contributed to animal husbandry and monitoring.

## Author Contributions

Conceptualization, S.M.M., A.S., J.P.L., D.A.H, L.J.R and A.R.P.; Methodology, S.M.M., O.H., A.S., S.K., A.C.P., D.P.S. and L.J.R.; Validation, K.S.O.; Formal Analysis, S.M.M., A.S., K.M.I., K.M.S. and A.C.P.; Investigation, S.M.M., O.H., K.S.O., A.S., S.K., C.J.S., A.C.W., G.W.M, and L.B.; Writing – Original Draft, S.M.M. and A.R.P.; Writing – Review & Editing, S.M.M., K.M.S., A.C.P., J.P.L, D.A.H and A.R.P.; Supervision, S.M.M., J.E.N, A.C.W.Z, and A.R.P.; Funding Acquisition, J.P.L, D.A.H., L.J.R. and A.R.P.

## Declaration of Interests

The authors declare no competing interests.

## STAR Methods

### Resource availability

#### Lead contact

Further information and requests for resources and reagents should be directed to and will be fulfilled by the lead contact, Allison Pettit.

#### Materials availability

This study did not generate new unique reagents.

#### Data and code availability

The datasets supporting the current study are available from the corresponding author on request.

### Experimental model and subject details

Animals were maintained under specific pathogen-free conditions in the Translational Research Institute Biological Resource Facility, fed standard chow with ad libitum water access and simulated diurnal cycle. Animal experiments were approved by The University of Queensland Health Sciences Ethics Committee and performed in accordance with the Australian Code of Practice for the Care and Use of Animals for Scientific Purposes.

#### Inbred strains and transgenic mice

C57BL/6J (wild type, WT) mice were obtained from Australian Resources Centre (Canning Vale, Western Australia, AU). Transgenic mouse lines were maintained in-house on C57BL/6J strain background; *Csf1r*-EGFP (RRID:IMSR_JAX:018549; (Sasmono et al., 2003)), *Siglec1*^Cre^ (RRID:IMSR_RBRC06239; (Karasawa et al., 2015)) and *Siglec1*^DTR^ (RRID:IMSR_RBRC04395; (Miyake et al., 2007)) sourced from Bio Resource Centre (Yokohama, Kanagawa, Japan), *R26*^ZsGreen^ (RRID:IMSR_JAX:007906; (Madisen et al., 2010)) originally sourced from Jackson Laboratory (Bar Harbor, Maine, USA). Experiments on naive mice were performed on adult mice (<6mth).

#### Macrophage depletion

For macrophage depletion experiments, data and samples from a previously published experiment (Batoon et al., 2019) were reanalysed. In brief, heterozygous adult male *Siglec1*^DTR^ mice (8wk old) were randomly allocated to groups treated with vehicle (0.9% sodium chloride) or 10 μg/kg body weight of diphtheria toxin (DT; MBL International Corporation, MA, USA) once daily via intra-peritoneal injection for up to 4 consecutive days. Tissues were harvested 24 hr after the last injection.

#### Emergency granulopoiesis

For stimulation of granulopoiesis, control (*Csf1r*-EGFP) and CD169-KO (homozygous *Siglec1*^DTR^) adult mice (<6mth) were injected twice daily subcutaneously, with vehicle (0.9% sodium chloride) or 125 μg/kg per injection recombinant human G-CSF (Filgrastim (Nivestim), Pfizer Australia) for up to 3 consecutive days.

### Method details

#### Tissue harvesting and processing

Mice were anesthetized and 1 mL blood collected into heparinised tubes by cardiac exsanguination. Femoral BM was collected for flow cytometry by flushing with a 5 mL syringe mounted with 27G needle and 2 x 5 mL of 2% fetal calf serum (FCS) in phosphate buffered saline (PBS). Spleens were dissected and weighed, and a portion of each spleen disaggregated using a either gentle MACS dissociator (Miltenyi Biotec, Cologne, Germany) (Jacobsen et al., 2014) or gently using a 1 mL plunger to push the tissue through a 70 μm strainer. The method of mechanical disruption did not impact the outcome. Lymph nodes (LN) were dissected and gently disaggregated through a 70 μm strainer using a 1 mL plunger. Dissociated cells were filtered through a 40-μm filter prior to antibody staining for flow cytometric analysis. Cell preparations from BM, spleen and LN were stained and analysed without red-blood-cell (RBC) lysis. Blood was subjected to RBC lysis with 10 volumes lysis buffer (150 mM NH_4_Cl, 10 mM NaHCO_3_, 1 mM EDTA) for 6 min at room temperature prior to washing and resuspension in 2% FCS/PBS for staining. For in situ analysis, tissues were fixed using 4% paraformaldehyde. Bones were decalcified using 14% EDTA for at least 2 weeks. Decalcified tissues were processed using a Tissue-Tek VIP 6 Processor (Sakura Finetek, Tokyo, Japan) and paraffin embedded. Spleens were infused with sucrose and cryo-embedded in OCT compound (Sakura Finetek, Tokyo, Japan).

#### Flow cytometry

Flow cytometry was performed on BM, spleen, lymph node and blood single cell suspensions prepared as described above. Cell counts were performed using a BC-5000 Vet Auto Haematology Analyser (Shenzhen Mindray Bio-Medical Electronics Co) to ensure a consistent number of cells was stained for each sample. When performed, enrichment of c-Kit^+^ (CD117) cells was achieved using magnetic activated cell sorting (Miltenyi Biotec) as described previously (Barbier et al., 2012). Cells were stained with antibody cocktails (as outlined in gating strategies and detailed in Key Resources Table), prepared in CD16/CD32 hybridoma 2.4G2 supernatant, for 30 min on ice and then washed with 2% FCS/PBS. Viability dyes (7-amino actinomycin D, Thermofisher, Massachusetts, USA; FVS700, BD Biosciences, New Jersey, USA) were used as per manufacturer’s instructions. Flow cytometric analyses were primarily performed using BD Fortessa flow cytometers. Data in Figure 6 was obtained using Cytoflex S and CyAn ADP Analyzers (Beckman Coulter, Brea, CA). Cell sorting was done using BD Aria Fusion sorters (HSPC sorts) or a MoFlo Astrios cell sorter (“macrophage” sorts). Data analysis was performed on singlet events and with dead cells excluded using FlowJo software version 10.7.1 (Tree Star Data Analysis Software; Tree Star, Ashland, OR). T-distributed stochastic neighbour embedding (t-SNE) data dimensionality reduction of the hematopoietic lineage stain was based on the following parameters: FSC, SSC, BV510 (CD11b), BV605 (CD115), BV650 (Sca1), BV785 (Ly6G), PerCPCy5.5 (CD11c), PECF594 (CD4/CD8), APC (Kit) and APCCy7 (B220).

#### Imaging flow cytometry

Cells were stained with antibody cocktails (as outlined in gating strategies and detailed in Key Resources Table), prepared in CD16/CD32 hybridoma 2.4G2 supernatant, for 30 min on ice and then washed with 2% FCS/PBS and resuspended at 20 million cells/mL for analysis on a three laser (405nm, 488nm, 642nm) Amnis Imagestream^X^ MkII Imaging Flow Cytometer (Luminex Corp, Austin, TX). Results were analysed using Amnis IDEAS Software v6.2.

#### Immunofluorescent staining and imaging

A Leica CM1950 cryostat (Leica Biosystems, Germany) was used to generate 5 μm sections of decalcified bone and spleen. Sections were stained with antibodies in blocking buffer (PBS containing 5% bovine serum albumin (BSA) and 0.1% Triton X-100) and washed by dipping in PBS. When required, sections were incubated with a PE-CF594 conjugated streptavidin secondary reagent in blocking buffer at room temperature for 1 hour. Sections were washed by dipping in PBS, and where indicated, sections were counterstained with 4′,6-diamidino-2-phenylindole (DAPI). Full antibody details are provided in the Key Resources Table. Bone sections in Figure 2 were stained with anti-F4/80-AF647 (BD Biosciences). Spleen sections in Figure 4 were stained with anti-B220-biotin (Biolegend), anti-F4/80-BV421 (BD Biosciences), anti-CD169-AF488 (Biolegend) and CD3-AF647 (Biolegend).

BM cell suspension was prepared as for flow cytometry, stained with anti-B220-AF594 (Biolegend) and anti-F4/80-BV421 (BD Biosciences), and cells imaged while in suspension. All imaging was conducted using an Olympus FV3000 confocal laser scanning microscope (Olympus, Tokyo, Japan). For morphometric analysis, digital microscopy images were analysed using ImageJ software as previously described (Inman et al., 2005). Colocalization analysis was performed using the Coloc 2 Image J plugin as described (Schindelin et al., 2012). Briefly, all images were coded and assessed blindly at least 3 sectional depths. Background intensity thresholds were applied using an ImageJ macro which measures pixel intensity across all immunostained and non-stained areas of the images and these were converted to percent staining area.

#### Immunohistochemical staining

Immunohistochemistry was performed on deparaffinised and rehydrated 4 μm sections as previously described (Batoon et al., 2019), using reagents documented in the Key Resources Table. Briefly, sections were blocked for endogenous peroxidase activity and subject to antigen retrieval by microwave in 10 mM sodium citrate (pH 6.0). Non-specific antibody binding was blocked using Background Sniper (Biocare Medical). Unconjugated anti-F4/80 was diluted in DaVinci Green Diluent (Biocare Medical) and incubated with sections for 90min. Sections were subsequently washed in Tris-buffered saline, incubated with biotinylated goat anti-rat secondary, washed again, and incubated with Vectastain Elite ABC-HRP, followed by development with ImmPACT DAB Peroxidase Substrate according to manufacturer’s instructions. Stained sections were imaged with an Olympus VS120 slide scanner.

#### Bulk RNA analysis of sorted cell populations

Cells populations for RNA analysis were sorted into 50% FCS/RPMI media. HSPC cell populations were subsequently washed with 2% FCS/PBS and cell pellets snap frozen for RNA extraction using QIAGEN RNAeasy mini kit as per manufacturer’s instructions. Reverse transcription (iScript cDNA synthesis kit) was performed prior to qPCR analysis with Taqman gene expression assays (see Key Resources Table). Assay reactions were run in Taqman Universal Master Mix II on a ViiA 7 Real-Time PCR System, and analysed with QuantStudio Real-Time PCR Software. RNA extraction from cell populations sorted by positive selection for macrophage markers was performed using Trizol reagent, followed by column clean-up using QIAGEN RNAeasy mini kit. The cDNA libraries for sequencing were generated as described previously (Bisht et al., 2019). The reverse transcription mastermix was prepared according to Picelli *et. al.* (Picelli et al., 2014) with the following modifications. SMARTScribe™ Reverse Transcriptase was used in place of SuperScript II, and the RNase inhibitor used was RNaseOUT. The template switching oligo did not contain any locked nucleic acids but was modified with biotin to prevent concatemerization. PCR pre-amplification was performed using 18 cycles. The amplicons were cleaned using Ampure beads as previously described (Picelli et al., 2014), and tagmented using the Nextera XT DNA sample preparation kit. Libraries were barcoded with Illumina adaptors from the Nextera XT 24-index kit and amplified with 13 cycles of PCR. Final libraries were subject to equimolar pooling at 2.5 nM. Libraries were gel purified to select for 200-600bp fragments and sequenced on an Illumina NextSeq500 instrument using the high output kit in the 75 bp paired-end configuration. Reads were mapped to the mouse genome (mm9) using Tophat2 with default parameters. Transcript abundance was computed with Cuffdiff2 (Trapnell et al., 2013). The MyGeneset databrowser (https://www.immgen.org/Databrowser19/DatabrowserPage.html) was used to compare transcript levels of the top 200 most abundant genes within each “macrophage” population across the ImmGen ULI-RNA-Seq datasets (GSE127267).

### Quantification and statistical analysis

Graphing and statistical analyses were conducted using GraphPad Prism 8.3.1, as documented in figure legends.

## Notes

### Competing Interest Statement

The authors have declared no competing interest.

