## Supplemental Figures and Tables for "Fragmentation of macrophages during isolation confounds analysis of single cell preparations from mouse hematopoietic tissues"

Figure S1. Gating Strategy for Delineating Murine Haematopoietic Cell Populations Applied to Spleen, Lymph Node and Peripheral Blood

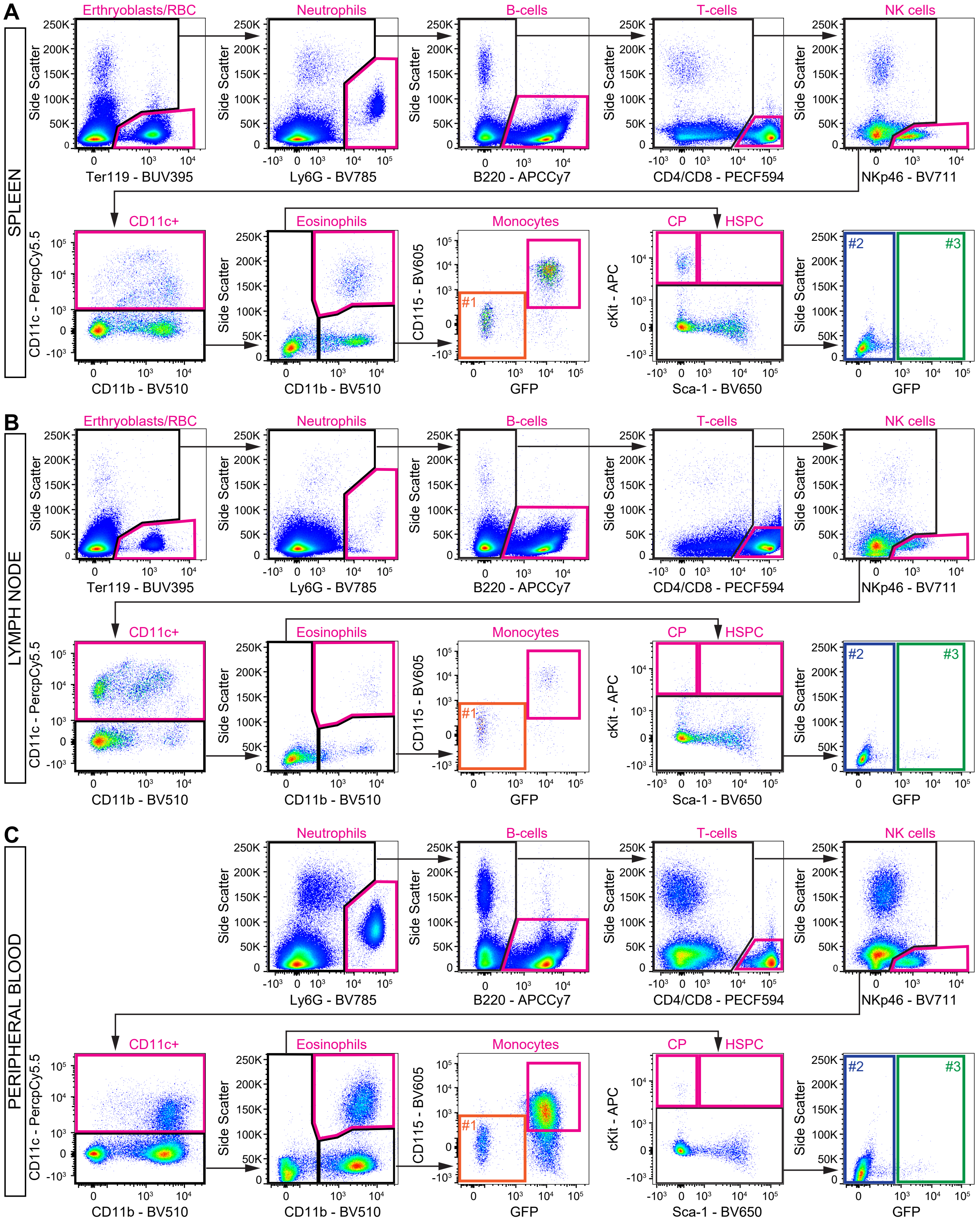

**Figure S1. Gating Strategy for Delineating Murine Hematopoietic Cell Populations Applied to Spleen, Lymph Node and Peripheral Blood.**

(A, B and C) Gating strategy employed to identify and exclude defined hematopoietic populations as a negative selection approach to identify putative macrophage populations in spleen (A), lymph node (B) and peripheral blood (C) from *Csf1r*-EGFP reporter mice. Excluded cell populations are labelled in magenta, and putative macrophage populations labelled in orange (#1), blue (#2) and green (3#). Cell frequency data for each population is summarized in Table S1.

Figure S2. Comparison of Macrophage Fidelity of Csf1r-EGFP and Siglec1Cre;R26ZsGreen Reporters in Bone Marrow Myeloid Populations by Imaging Flow Cytometry.

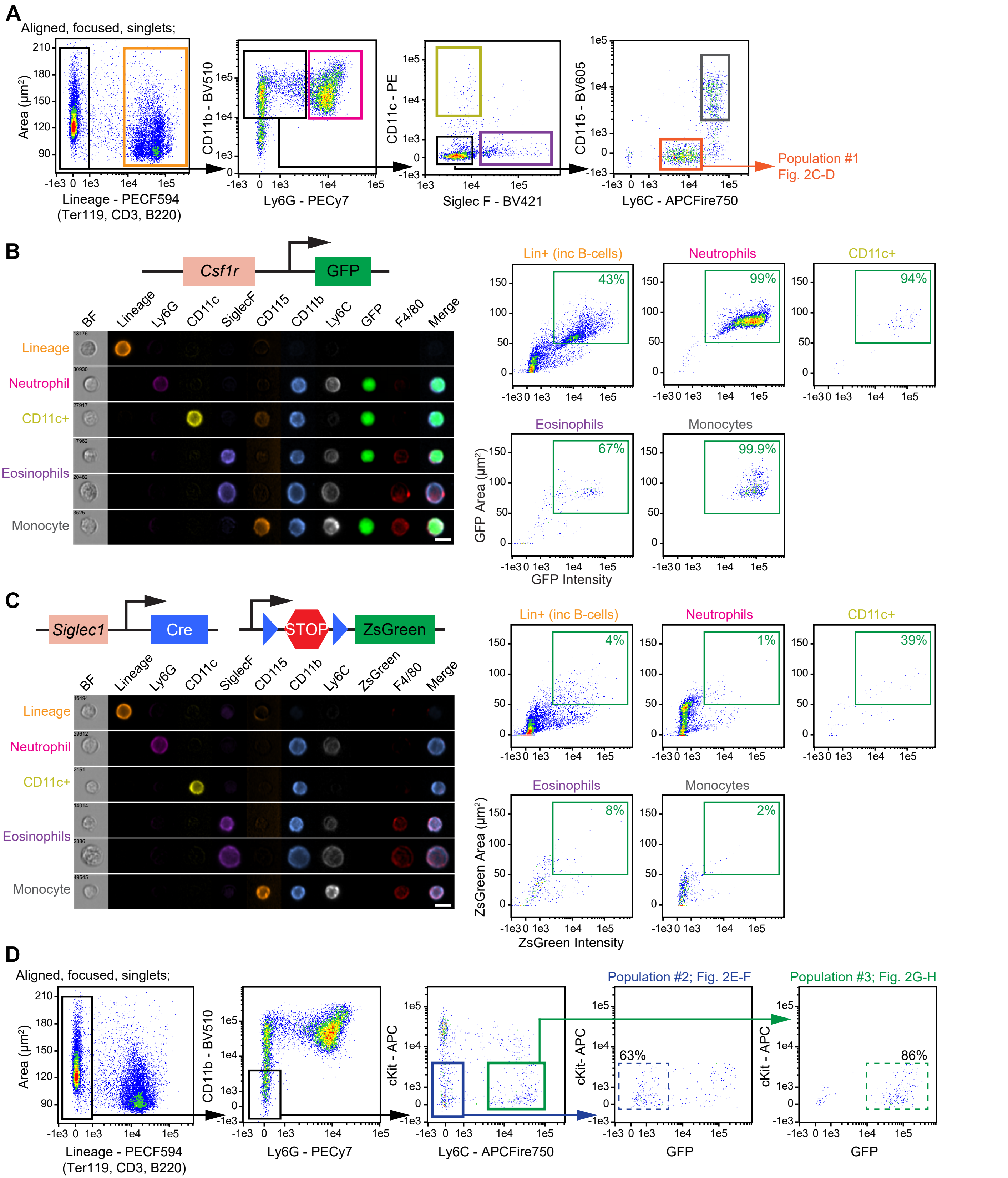

**Figure S2. Comparison of Macrophage Fidelity of *Csf1r*-EGFP and *Siglec1*$^{Cre}$;*R26*$^{ZsGreen}$ Reporters in Bone Marrow Myeloid Populations by Imaging Flow Cytometry.**

(A) Imaging flow cytometry gating strategy employed to identify CD11b$^{+}$ myeloid populations in bone marrow. The gate colours correspond to cell populations as follows: Light orange, Lineage$^{+}$. Magenta, neutrophils. Yellow, CD11c$^{+}$. Purple, eosinophils. Grey, monocytes. Dark orange, Population #1. Representative images of Population #1 are displayed in Figure 2C and 2D.

(B) GFP reporter expression in *Csf1r*-EGFP mice was documented within non-macrophage CD11b$^{+}$ myeloid cells with representative images and dot plot quantitation. The merge images consist of GFP reporter, CD11b and F4/80. Scale bar, 10 µm.

(C) ZsGreen reporter expression in *Siglec1*$^{Cre}$;*R26*$^{ZsGreen}$ mice was absent from non-macrophage CD11b$^{+}$ myeloid cells as documented with representative images and dot plot quantitation. The merge images consist of ZsGreen reporter, CD11b and F4/80. Scale bar, 10 µm.

(D) The gating strategy used to delineate the putative macrophage populations #2 (representative images displayed in Figure 2E and 2F) and #3 (representative images displayed in Figure 2G and 2H) is outlined in the first three panels. The following two panels document that in *Csf1r*-EGFP mice the Lineage$^{-}$Kit$^{-/lo}$Ly6C$^{-}$ population is predominantly GFP$^{-}$ and the Lineage$^{-}$Kit$^{-/lo}$Ly6C$^{+}$ population is predominantly GFP$^{+}$.

Figure S3. Imaging Flow Cytometry Approach Used to Interrogate Macrophage Marker Staining on Mature Haematopoietic Lineages in BM and Spleen

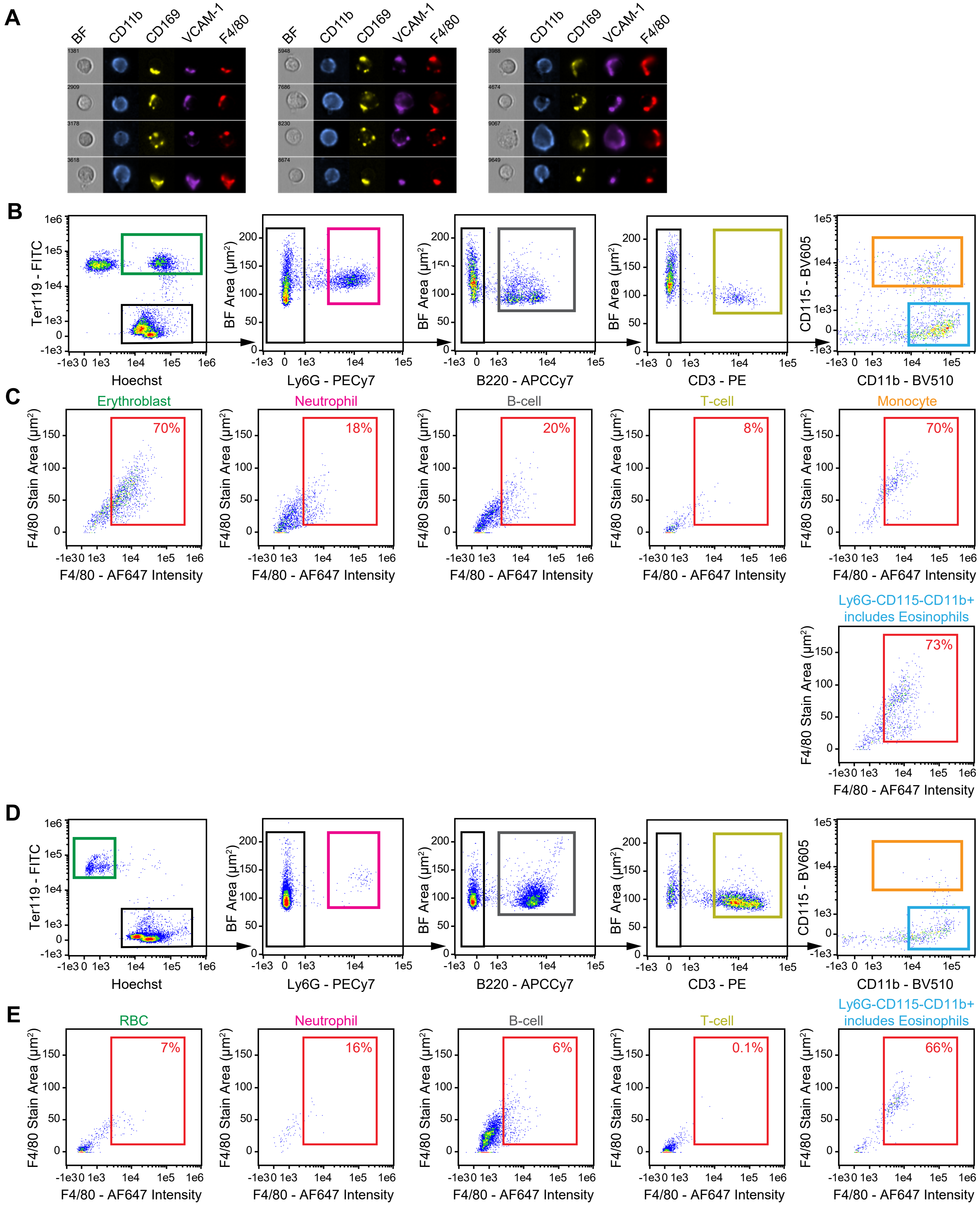

**Figure S3. Imaging Flow Cytometry Approach Used to Interrogate Macrophage Marker Staining on Mature Hematopoietic Lineages in BM and Spleen.**

(A) Imaging flow cytometry was used to visualize $CD11b^{+}CD169^{+}VCAM\text{-}1^{+}F4/80^{+}$ events in BM isolated from C57BL/6J mice. Representative images demonstrate that staining distribution for CD169, VCAM-1 and F4/80 was coincident in a restricted polarized pattern, which contrasted with the cell surface expression of CD11b. Scale bar, 10 µm.

(B) Imaging flow cytometry gating strategy used identify mature hematopoietic lineages in BM, as visualized in Figure 3A. The gate colours correspond to cell populations as follows: Green, $Ter119^{+}Hoechst^{+}$ erythroblasts. Magenta, $Ly6G^{+}$ neutrophils. Grey, $B220^{+}$ B-cells. Yellow, $CD3\varepsilon^{+}$ T-cells. Orange, $CD115^{+}$ monocytes. Blue, $Ly6G^{-}CD115^{-}CD11b^{+}$ cells including neutrophil precursors and eosinophils.

(C) Frequency of F4/80+ events for each of the populations gated in (B). For F4/80+ events, the stain area for F4/80 and a defining cell surface marker is graphed in Figure 3C.

(D) Imaging flow cytometry gating strategy used identify mature hematopoietic lineages in spleen, as visualized in Figure 3B.

(E) Frequency of F4/80+ events for each of the populations gated in (D). Where sufficient F4/80+ events were imaged, the stain area for F4/80 and a defining cell surface marker is graphed in Figure 3D. Note that the sensitivity for detecting $F4/80^{+}$ events by imaging flow cytometry is substantially less with F4/80-AF647 than F4/80-BV421.

Figure S4. Conventional Flow Cytometry Macrophage Marker Staining Profiles Across Hematopoietic Cell Types and Tissues

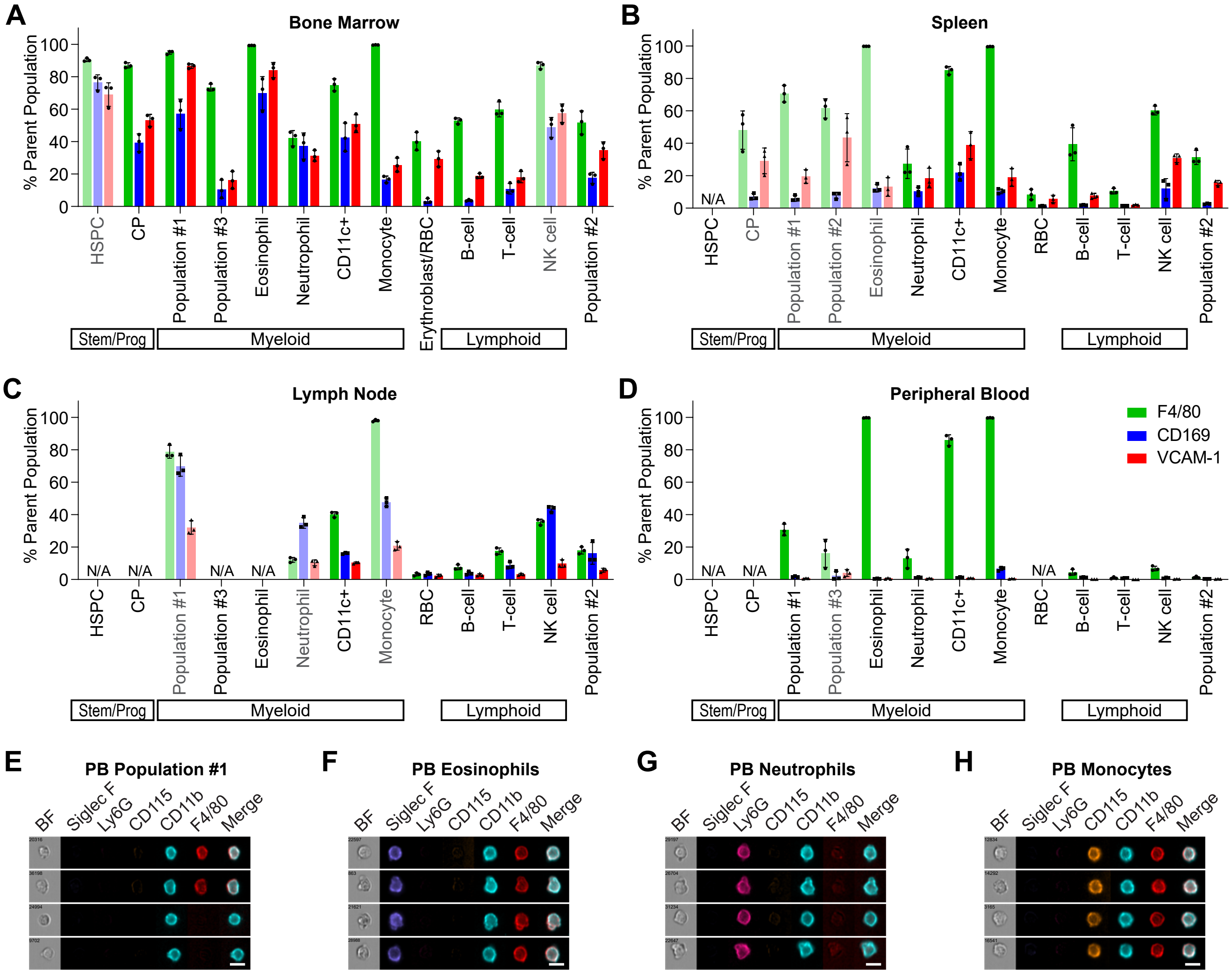

**Figure S4. Conventional Flow Cytometry Macrophage Marker Staining Profiles Across Hematopoietic Cell Types and Tissues**

(A – D) Frequency of F4/80+ (green bars), CD169+ (blue bars) and VCAM-1+ (red bars) event assessed within each of the hematopoietic cell populations delineated in Figure 1A and Figure S1 for bone marrow (A), spleen (B), lymph node (C) and peripheral blood (D). For cell populations that represent <0.5% live cells data have been presented as semi-transparent bars, and for cell populations <0.05% live cells data are not presented. N/A, not applicable. Data are represented as mean ± s.d.*;* *n* = 3.

(E – H) Imaging flow cytometry visualization of peripheral blood (PB) cells: population #1 (E), eosinophils (F), neutrophils (G) and monocytes (H). Cell surface F4/80 expression was confirmed within all of these populations with the exception of neutrophils.

Figure S5. Hematopoietic Stem/Progenitor Cell Gating and Sorting BM Populations by Positive Selection for Macrophages mMarkers

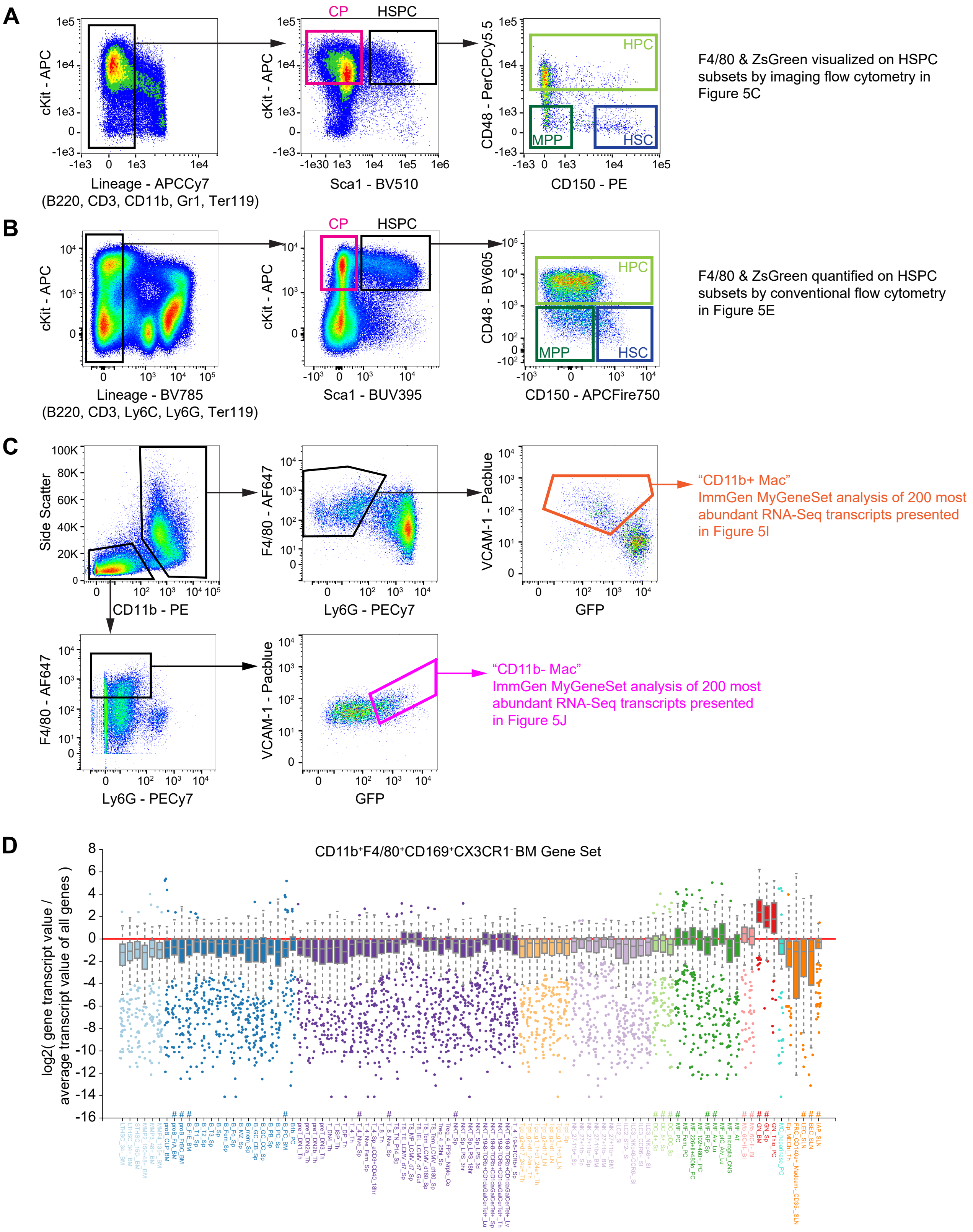

**Figure S5. Hematopoietic Stem/Progenitor Cell Gating and Sorting BM Populations by Positive Selection for Macrophages Markers**

(A and B) Gating strategy employed to identify hematopoietic stem and progenitor populations in Kit-enriched bone marrow from *Siglec1*$^{Cre}$;*R26*$^{ZsGreen}$ mice by imaging flow cytometry (A and Figure 5C) and conventional flow cytometry (B and Figure 5E). Committed progenitors (CP) were Lin$^{-}$cKit$^{+}$Sca1$^{-}$. Lin$^{-}$cKit$^{+}$Sca1$^{+}$ hematopoietic stem and progenitor cells (HSPC) were subsetted into: CD48$^{+}$ hematopoietic progenitor cell (HPC), CD48$^{-}$CD150$^{-}$ multipotent progenitor (MPP) and CD48$^{-}$CD150$^{+}$ hematopoietic stem cell (HSC).

(C) Positive selection strategy employed to sort “macrophage” populations from BM of *Csf1r*-EGFP mice for RNA-Seq analysis, as presented in Figure 5I and 5J.

(D) The top 200 most abundant transcripts, as determined by RNA-Seq, from a CD11b$^{+}$F4/80$^{+}$CD169$^{+}$CX3CR1$^{-}$ sorted BM population from C57BL/6J mice were assessed using the ImmGen MyGeneSet analysis tool. Prevalence of these transcripts across the entire the ImmGen ULI RNA-seq data-set (GSE127267) is displayed. The transcript list was highly represented in ImmGen granulocyte populations (red bars). # marks the ImmGen ULI RNA-seq populations that were included in the analysis presented in Figure 5I and 5J.

Figure S6. Transcript Abundance of Macrophage, Neutrophil and Erythroblastic Markers in Published BM Monocyte and Macrophage RNA-Seq Data.

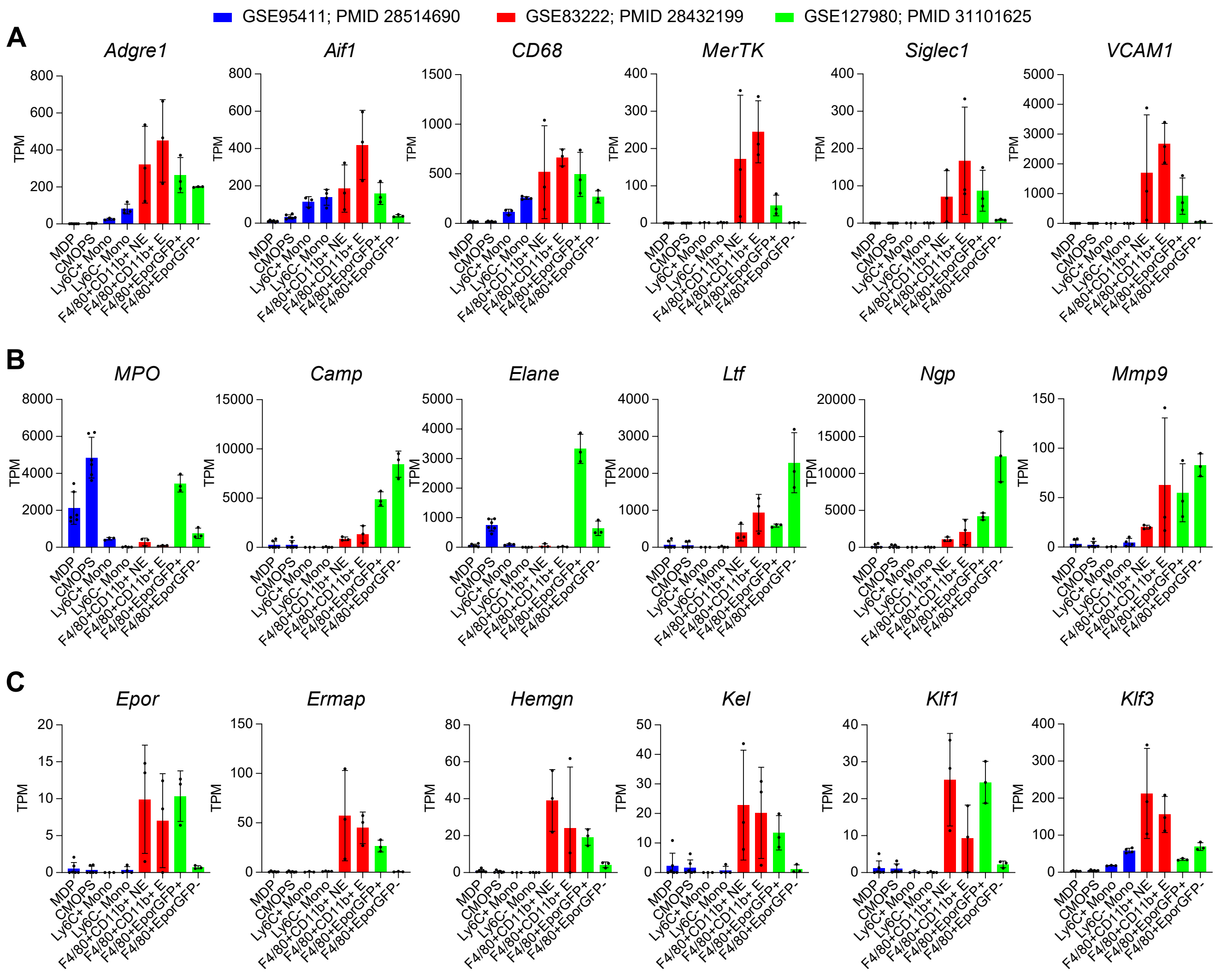

**Figure S6. Transcript Abundance of Macrophage, Neutrophil and Erythroblastic Markers in Published BM Monocyte and Macrophage RNA-Seq Data.**

(A – C) Comprehensive meta-analysis of RNA-Seq data from the mouse mononuclear phagocyte system (Summers et al., 2020) included bone marrow monocyte-macrophage samples from three studies (NCBI-GEO Series; GSE95411, GSE83222, GSE127980). Transcript abundance data for (A) monocyte-macrophage, (B) neutrophil and (C) erythroblastic markers from these studies is displayed. TPM, transcripts per million. MDP, macrophage-dendritic cell progenitor. cMoP, monocyte-committed common monocyte progenitor. Mono, monocyte. NE, non-engulfing (ie non-phagocytic). E, engulfing (ie phagocytic). Data are represented as mean ± s.d.; n = 3.

Figure S7. CD169 Deletion is Permissive of G-CSF Stimulated Granulopoiesis in the BM, but Blunts Egress of Neutrophils into Peripheral Circulation.

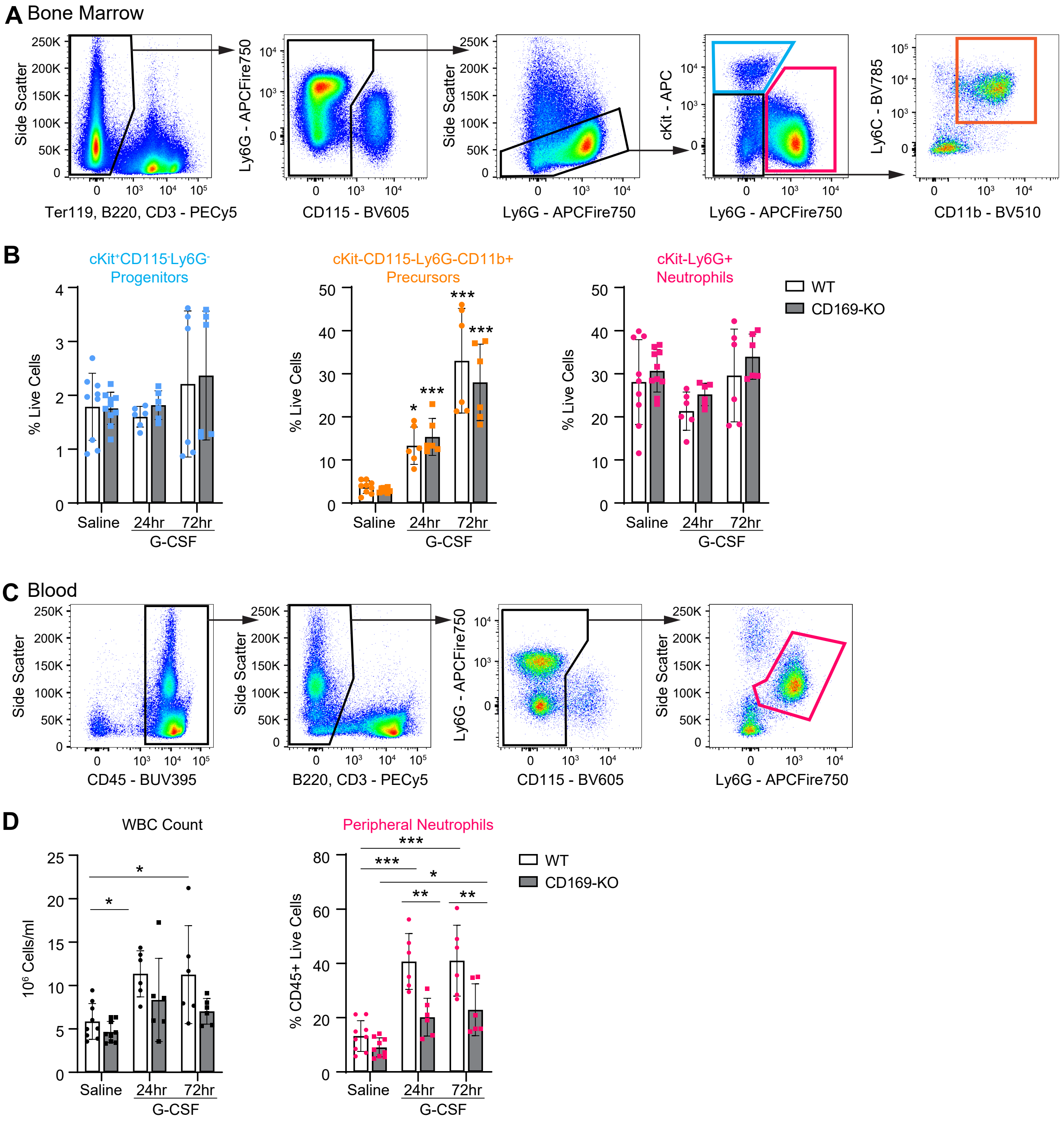

**Figure S7. CD169 Deletion is Permissive of G-CSF Stimulated Granulopoiesis in the BM, but Blunts Egress of Neutrophils into Peripheral Circulation.**

(A) Conventional flow cytometry gating strategy employed to examine G-CSF stimulated emergency granulopoiesis in the bone marrow, shown for a representative saline treated control. Blue gate, $Lin^-$ $cKit^+CD115^-Ly6G^-$ progenitors. Orange gate, $Lin^-cKit^-CD115^-Ly6G^-CD11b^+$ precursors. Pink gate, cKit- $Ly6G^+$ neutrophils.

(B) The proportion of these three populations in the bone marrow following bi-daily 125 µg/kg G-CSF injections is displayed. No statistical differences were observed between wild-type (WT) and CD169-KO mice for any population at any time-point. Data are pooled from 4 independent experiments and represented as mean ± s.d.; $n$ = 6-9. Two-way ANOVA with Tukey correction for multiple comparisons. Statistical differences from saline-treated control are indicated within each genotype. *p = 0.012, ***p < 0.0001.

(C) Conventional flow cytometry gating strategy employed to quantify peripheral neutrophils, shown for a representative saline treated control. Pink gate, $Ly6G^+$ neutrophils.

(D) Bi-daily 125 µg/kg G-CSF injections yields an increase in both peripheral blood cell count and the proportion of $CD45^+$ blood cells that are neutrophils in wild-type (WT) mice. G-CSF response to both these parameters are blunted in CD169-KO mice. Data are pooled from 4 independent experiments and represented as mean ± s.d.; $n$ = 6-9. Two-way ANOVA with Tukey correction for multiple comparisons. *p < 0.05, **p < 0.01. *** p<0.0001. WBC, white blood cell count.

**Table S1: Frequency of hematopoietic cell populations in single cell tissue preparations from *Csf1r*-EGFP mice.**

n = 3 adult (<6mth) male *Csf1r*-EGFP mice

| | Bone Marrow | Spleen | Lymph Node | Blood |
|---|---|---|---|---|
| Erythroblast/RBC | 34 ± 7 | 32 ± 10 | 2.4 ± 0.2 | N/A (RBC lysis) |
| Neutrophils | 29 ± 5 | 1.0 ± 0.1 | 0.15 ± 0.06 | 9 ± 3 |
| B-cells | 17 ± 1 | 44 ± 11 | 41 ± 4 | 61 ± 7 |
| T-cells | 1.1 ± 0.1 | 17 ± 2 | 53 ± 3 | 12 ± 2 |
| NK cells | 0.43 ± 0.04 | 1.4 ± 0.2 | 0.38 ± 0.07 | 2.4 ± 0.6 |
| CD11c+ Cells | 0.60 ± 0.05 | 0.6 ± 0.1 | 1.1 ± 0.3 | 1.3 ± 0.6 |
| Eosinophils | 2.6 ± 0.3 | 0.29 ± 0.05 | 0.04 ± 0.01 | 2.2 ± 0.6 |
| Monocytes | 4.2 ± 0.6 | 0.8 ± 0.3 | 0.07 ± 0.03 | 5 ± 3 |
| Committed Prog | 1.3 ± 0.1 | 0.14 ± 0.006 | 0.008 ± 0.003 | 0.020 ± 0.009 |
| HSPC | 0.19 ± 0.03 | 0.010 ± 0.005 | N/A | N/A |
| **Sum defined cell populations:** | **91 ± 1** | **97 ± 2** | **98 ± 0.4** | **92 ± 1** |
| Population #1 | 5.5 ± 1.4 | 0.25 ± 0.07 | 0.13 ± 0.03 | 0.79 ± 0.30 |
| Population #2; GFP- | 1.6 ± 0.2 | 2.2 ± 1.3 | 1.5 ± 0.4 | 5.8 ± 1.2 |
| Population #3; GFP+ | 0.90 ± 0.05 | 0.18 ± 0.18 | 0.022 ± 0.006 | 0.10 ± 0.12 |
| **Sum (Pop#1-3):** | **8.0 ± 1.3** | **2.7 ± 1.6** | **1.7 ± 0.4** | |

**Table S2: Frequency of F4/80⁺ staining on putative macrophage populations.**

Greyed out cells indicate that too few events were counted to provide reliably enumeration.

| | Bone Marrow | Spleen | Lymph Node | Blood |
|---|---|---|---|---|
| Population #1 | 95 ± 1 | 71 ± 5 | 79 ± 4 | 31 ± 6 |
| Population #2; GFP- | 52 ± 7 | 31 ± 4 | 18 ± 2 | 1 ± 1 |
| Population #3; GFP+ | 73 ± 2 | 62 ± 6 | | |

**Table S3: Frequency of hematopoietic cell populations in bone marrow from WT and CD169-KO mice.**

n = 3 adult (<6mth) female mice per genotype

| | C57Bl/6 | CD169-KO |
|---|---|---|
| Erythroblast/RBC | 40 ± 3 | 41 ± 4 |
| Neutrophils | 20 ± 2 | 21 ± 2 |
| B-cells | 22 ± 1 | 19 ± 3 |
| T-cells | 1.7 ± 0.4 | 1.7 ± 0.8 |
| NK cells | 0.33 ± 0.14 | 0.35 ± 0.11 |
| CD11c+ Cells | 0.34 ± 0.02 | 0.31 ± 0.03 |
| Eosinophils | 3.3 ± 0.3 | 4.0 ± 0.5 |
| Monocytes | 1.9 ± 0.8 | 2.4 ± 1.0 |
| Committed Prog | 1.3 ± 0.1 | 1.4 ± 0.1 |
| HSPC | 0.22 ± 0.03 | 0.21 ± 0.03 |
| **Sum defined cell populations:** | **91 ± 1** | **92 ± 2** |
| Population #1 | 4.6 ± 0.4 | 4.7 ± 1.5 |
| Population #2; Ly6C- | 1.5 ± 0.1 | 1.4 ± 0.2 |
| Population #3; Ly6C+ | 1.44 ± 0.05 | 1.39 ± 0.03 |
| **Sum (Pop#1-3):** | **7.5 ± 0.5** | **7.5 ± 1.6** |
